# Ovoselenol, a Selenium-containing Antioxidant Derived from Convergent Evolution

**DOI:** 10.1101/2024.04.10.588772

**Authors:** Chase M. Kayrouz, Kendra A. Ireland, Vanessa Ying, Katherine M. Davis, Mohammad R. Seyedsayamdost

**Author notes:** These authors contributed equally (Chase M. Kayrouz, Kendra A. Ireland).

## Abstract

Selenium is an essential micronutrient, but its presence in biology has been limited to protein and nucleic acid biopolymers. The recent identification of the first biosynthetic pathway for selenium-containing small molecules suggests that there is a larger family of selenometabolites that remains to be discovered. Using a bioinformatic search strategy that relies on mapping of composite active site motifs, we identify a recently evolved branch of abundant and uncharacterized metalloenzymes that we predict are involved in selenometabolite biosynthesis. Biochemical studies confirm this prediction and show that these enzymes form an unusual C–Se bond onto histidine, thus giving rise to a novel selenometabolite and potent antioxidant that we have termed ovoselenol. Aside from providing insights into the evolution of this enzyme class and the structural basis of C–Se bond formation, our work offers a blueprint for charting the microbial selenometabolome in the future.

## Main Text

Low-molecular-weight (LMW) selenols are an emerging and intriguing class of biomolecules. They likely possess biological functions similar to the chemically-related LMW thiols, whose versatile reactivities and ability to access several oxidation states allow them to play a central role in myriad biological processes^1,2^. However, unlike the LMW thiol family, which boasts many characterized members in both primary and secondary metabolism, the list of known LMW selenols is exceedingly short, with the current compendium derived from Se-specific biosynthetic pathways comprising only selenosugars and selenoneine^3,4^ (**Fig. 1a**). Both were recently discovered through a selenium-focused genome mining effort, which not only revealed a new biosynthetic route but also pointed to an uncharted chemical space of natural selenometabolites.

**Figure 1.**
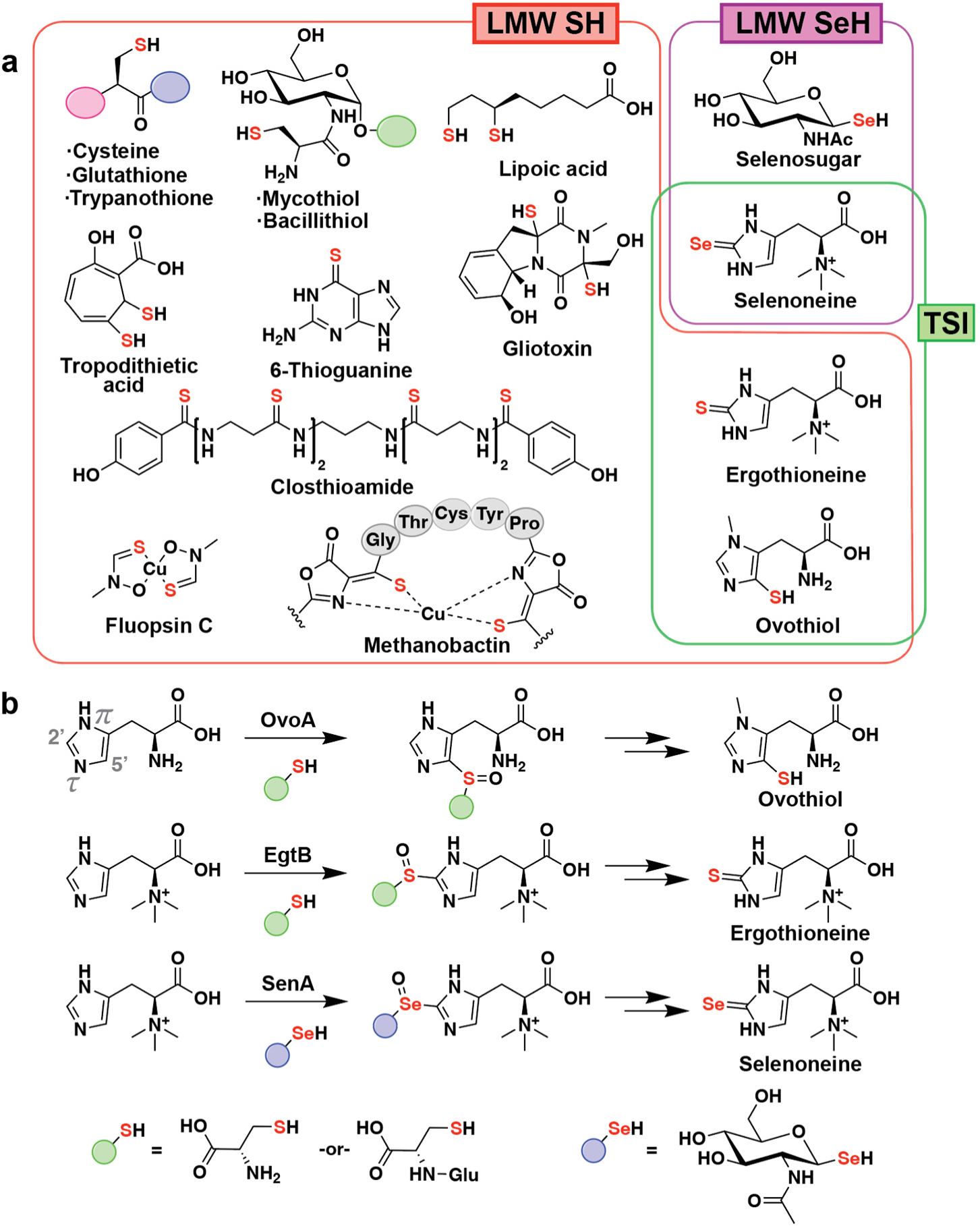
LMW thiols and selenols in biology. (**a**) Select LMW thiols (SH), all known LMW selenols (SeH) formed by specific biosynthetic pathways, and the thio/selenoimidazole (TSI) family. LMW thiols are represented in primary and secondary metabolism, with tropodithietic acid, 6-thioguanine, gliotoxin, closthioamide, fluopsin C, and methanobactin serving as members of the latter group. (**b**) NHISS-catalyzed C–S/Se bond forming reactions involved in the biosynthesis of TSIs.

A number of approaches can be envisioned for exploring this new chemical space. One potentially fruitful strategy revolves around the thio/selenoimidazoles (TSIs), a small but enticing class of thiol/selenol-functionalized histidine derivatives in biology, consisting of ergothioneine, ovothiol, and selenoneine^5^ (**Fig. 1a**). Typically synthesized by bacteria and fungi, these LMW thiols/selenols are thought to play important, redox-based roles in the lives of their producers, as well as in higher organisms for which TSIs are diet-derived. This includes humans, which accumulate high concentrations of ergothioneine in various tissues through uptake by the membrane transporter OCTN1^6,7^. TSIs are also interesting from a biosynthetic perspective, as they involve a unique metalloenzyme class that we have termed non-heme iron sulfoxide/selenoxide synthases (NHISS) responsible for forging C–S/Se bonds. In ovothiol and ergothioneine biosynthesis, the NHISS homologs OvoA and EgtB, respectively, catalyze oxidative C–S bond formation between cysteine and histidine derivatives *en route* to the final products^8,9^ (**Fig. 1b, Supplementary Fig. 1**). The recently discovered NHISS SenA carries out an analogous reaction using a selenosugar as a selenium source, in place of cysteine, to generate a selenoxide intermediate in the selenoneine biosynthetic pathway^4^ (**Fig. 1b, Supplementary Fig. 1**).

The small number of characterized NHISS enzymes led us to wonder whether divergent homologs have involved in the biosynthesis of yet-unknown selenometabolites . To approach this question, we compiled the active site motifs of previously characterized family members and used these to traverse the phylogenetic tree of the NHISS family in search of homologs with divergent substrate binding residues. This analysis revealed a large and previously uncharacterized clade of NHISS enzymes, which we predicted may carry out new chemistry. Using a combination of biochemical and structural biology techniques, we show they are involved in the production of a novel selenometabolite that we term ovoselenol. Aside from uncovering its complete biosynthetic pathway and providing insights into the structure and evolution of NHISS enzymes, our results reveal a previously unknown chemotype in the TSI family and highlight that the microbial selenometabolome is now ripe for exploration.

### Phylogenetic analysis of the NHISS enzyme family

We began with the hypothesis that members of the NHISS family that catalyze new reactions would have active site motifs divergent from those of previously characterized members. To date, five subfamilies of bacterial NHISS enzymes have been studied. Type I, II, and IV EgtBs (hereafter referred to as EgtB-I, EgtB-II, and EgtB-IV) catalyze C–S bond formation between cysteine and the imidazole 2′-carbon of hercynine (*N*α-trimethylhistidine) during ergothioneine biosynthesis^9-11^ (**Fig. 1b, Supplementary Fig. 1**). OvoA, representing a fourth subfamily, introduces a C–S bond between cysteine and the 5′-carbon of the histidine-imidazole^8^. Finally, the recently identified subfamily containing SenA catalyzes C–Se bond formation between a selenosugar and the 2′-carbon of hercynine^4^. Type III EgtBs are of fungal origin and will not be discussed further here^10,12^. Crystal structures of EgtB-I, EgtB-II, OvoA, and SenA have been reported, revealing key active site residues involved in metal binding, substrate positioning, and catalysis^10,13-15^ (**Fig. 2a-d**). Examination of these residues by multiple sequence alignment appears to be sufficient to predict the subfamily of known NHISS enzymes. We therefore reasoned that we could compress any NHISS protein sequence into a short, composite motif that would be predictive of its function (**Fig. 2e**). Applying this strategy, we retrieved all NHISS homologs from the NCBI database, extracted the relevant residues for each member by multiple sequence alignment, and mapped these composite motifs onto a phylogenetic tree of the retrieved protein sequences. A subset of this tree is displayed (**Fig. 2f, Supplementary Fig. 2**) and illustrates the family’s evolutionary trajectory, highlighting an important early bifurcation between the OvoA-like enzymes (OvoA and EgtB-IV) and the EgtB-like enzymes (EgtB-I, EgtB-II, and SenA). This analysis indicates that the vast majority of members fall into one of the previously characterized classes (**Supplementary Table 4**), as judged by their composite motifs. However, closer examination reveals a new clade, diverging most recently from the SenA family, which bears a unique composite motif and has yet to be characterized (**Fig. 2e**, ‘Unknown NHISS’). We therefore predicted that this divergent clade, a sixth class of NHISS, may be responsible for new chemistry, possibly involving selenium given its close homology to the SenA subfamily.

**Figure 2.**
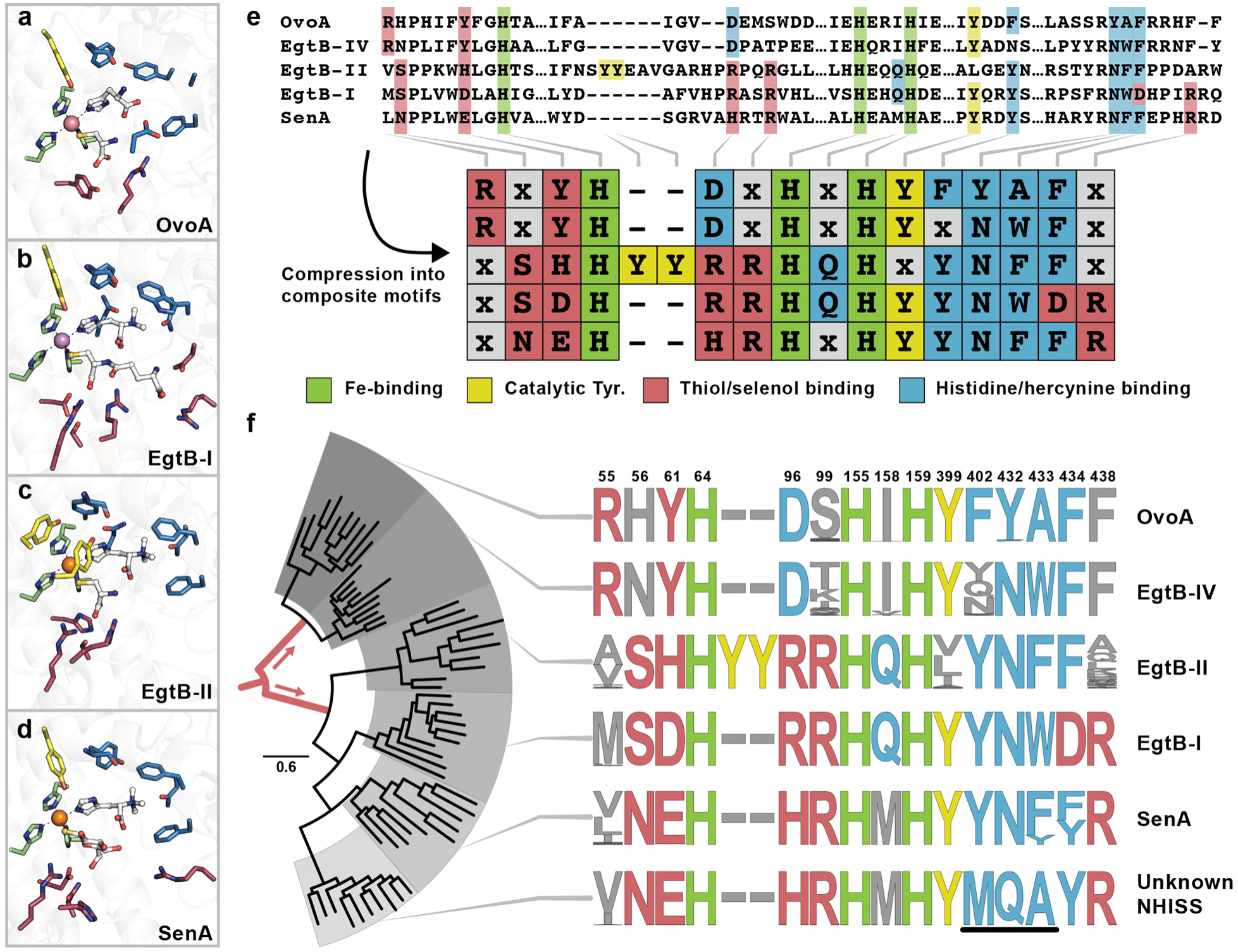
Identification of divergent NHISS clades with potentially novel functions. (**a-d**) Active site architectures of NHISS subfamilies, with important residues colored according to function (PDB accession codes 8KHQ^14^, 4X8D^13^, 6O6L^10^, 8K5I^15^, respectively); see panel **e** for a legend. (**e**) Compression of full-length multiple sequence alignments into composite motifs, which encapsulate the relevant residues, colored according to panels **a**-**d**. (**f**) Mapping the composite motifs onto a phylogenetic tree of the NHISS family reveals a new clade that features a unique composite motif and diverges recently from the SenA subfamily. Tree root is highlighted in red, representing the family’s initial bifurcation. Residue position numbering for OvoA from *Hydrogenimonas thermophila* is shown. The key residues distinguishing the unknown NHISS from the other subfamilies are underlined.

### A widespread BGC containing a divergent NHISS encodes the biosynthesis of ovoselenol

Within the current sequence database, the new NHISS subfamily contains 228 members derived almost exclusively from aquatic γ-proteobacteria (**Fig. 3a, b**). To elucidate their function, we began by analyzing their genomic contexts to pinpoint surrounding genes that may be involved in the same biosynthetic pathway. Interestingly, many look conspicuously similar to the selenoneine (*sen*) biosynthetic gene clusters (BGCs), featuring a selenosugar synthesis cassette (*senBC*) in addition to the divergent NHISS. An additional fourth gene, a close homolog of the ovothiol *N*π-methyltransferase (**Supplementary Fig. 1**) that is not present in *sen* clusters, is also often co-localized with the new NHISS. These four genes consistently co-occur, albeit separated chromosomally in some cases (**Fig. 3a, Supplementary Fig. 3**). Furthermore, the majority of these organisms lack the histidine *N*α-trimethylase gene, *egtD*, suggesting a substrate different from that of the SenA or EgtB subfamilies. We therefore speculated that these divergent NHISS enzymes may be catalyzing C–Se bond formation between histidine and a selenosugar, followed by imidazole methylation by the associated methyltransferase.

**Figure 3.**
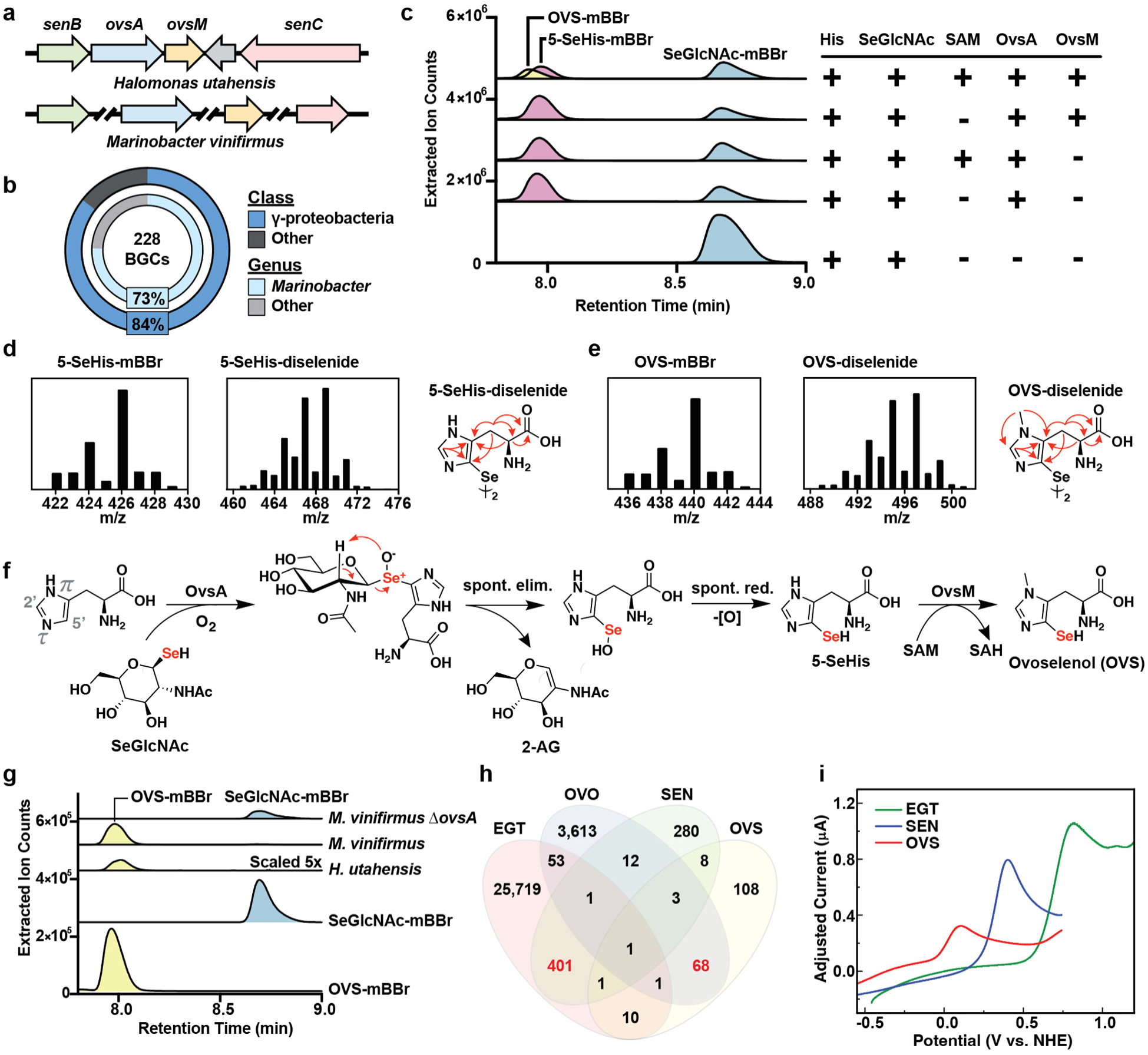
Discovery of ovoselenol (OVS) and its biosynthetic pathway. (**a**) Genes *ovsA* (NHISS) and *ovsM* (methyltransferase) are clustered with selenosugar biosynthesis genes *senB/C* in *Halomonas utahensis.* The gray arrow represents a hypothetical gene not conserved among *ovs* clusters. In other strains, such as *Marinobacter vinifirmus*, the genes co-occur but are chromosomally separated. (**b**) *Ovs* genes are found predominantly in *Marinobacter* and other γ-proteobacteria species. (**c**) *In vitro* reconstitution of OVS biosynthesis reveals 5′-selenylation of histidine by recombinant OvsA in the presence of SeGlcNAc, followed by SAM-dependent Nπ-methylation by OvsM. Extracted ion chromatograms of mBBr-derivatives of SeGlcNAc, 5-selenohistidine (5-SeHis), and OVS are shown. (**d, e**) Mass spectra of mBBr-derivatized and underivatized 5-SeHis and OVS. Relevant ^1^H-^13^C HMBC NMR correlations (red arrows) used to solve the structure are shown. (**f**) Complete biosynthetic pathway of OVS. See text for details. (**g**) Identification of mBBr-derivatized OVS in crude extracts of wild type *H. utahensis* and *M. vinifirmus*. Genetic deletion of *ovsA* in the latter results in a mutant that fails to produce OVS, accumulating SeGlcNAc as a result. Extracted ion chromatograms of mBBr-derivatives of SeGlcNAc and OVS are shown, alongside mBBr derivatives of pure standards. (**h**) Co-occurrence of BGCs for ergothioneine (EGT), ovothiol (OVO), selenoneine (SEN), and OVS in all sequenced microbes. Sequences from undefined taxa were omitted from this analysis. The combinations of EGT/SEN and OVO/OVS (shown in red) are disproportionally overrepresented. (**i**) Peak oxidation potentials of OVS, and of EGT and SEN for comparison, measured by cyclic voltammetry. See text and SI for details.

To test this hypothesis, we recombinantly expressed and purified the novel NHISS enzyme and its adjacent methyltransferase from *Halomonas utahensis*. Upon joint incubation of these two enzymes with the Se-donor from the *sen* pathway, *N*-acetyl-1-seleno-β-D-glucosamine (SeGlcNAc), and *S*-adenosylmethionine (SAM), the reactions were quenched with the thiol/selenol-derivatization reagent monobromobimane (mBBr) and analyzed by high-performance liquid chromatography-coupled mass spectrometry (HPLC-MS) (**Fig. 3c**). Remarkably, reaction of the novel NHISS with SeGlcNAc and histidine resulted in rapid formation of a new species with a single selenium atom, as judged by the characteristic MS isotope pattern of selenium-containing compounds (**Fig. 3d**). This intermediate was converted to a product 14 Da heavier upon the addition of the methyltransferase and SAM, indicating a single methyl transfer (**Fig. 3e**). Purification and NMR-based structure elucidation of both underivatized species revealed 5-selenohistidine and *N*π-methyl-5-selenohistidine as products of the NHISS and the methyltransferase, respectively (**Fig. 3d, e, Supplementary Tables 5 and 6, Supplementary Figs. 7 and 8**). The latter is the selenium isologue of ovothiol, a never before observed chemical species, which we now term ovoselenol. Consequently, we designate these divergent NHISS enzymes and their associated methyltransferases as OvsA and OvsM, respectively.

Additional assays confirmed that OvsA does not accept Nπ-methylhistidine nor does OvsM methylate histidine, suggesting a clear order of biosynthetic events consisting of histidine selenylation followed by *N*π-methylation (**Fig. 3f** and **Supplementary Fig. 4**). Analogous to SenA reactivity^4^, selenylation by OvsA likely proceeds through enzyme-catalyzed, oxidative C–Se bond formation to furnish a short-lived selenoxide that rapidly eliminates to release the sugar moiety in the form of 2-acetamidoglucal (2-AG), a species that we observe in the full reaction mixture (**Fig. 3f** and **Supplementary Fig. 5**).

We next sought to demonstrate that ovoselenol is the intended product of the *ovs* pathway by analyzing the metabolites produced by organisms harboring the *ovs* BGC (**Fig. 3a, g**). Indeed, we observed clear production of ovoselenol upon culturing *H. utahensis* and the genetically tractable bacterium *Marinobacter vinifirmus* in the presence of sodium selenite. Furthermore, targeted gene disruption of *ovsA* in *M. vinifirmus* resulted in a mutant (*ΔovsA*) unable to produce ovoselenol (**Fig. 3g, Supplementary Fig. 6**). This mutant instead accumulated SeGlcNAc, confirming the identity of the selenosugar substrate and providing the first direct evidence for its production in bacteria. Together, these results establish ovoselenol as a new selenometabolite generated by the *ovs* BGC.

With ovothiol, ergothioneine, selenoniene, and now ovoselenol, the TSI family encompasses four distinct chemotypes. Analysis of the frequency of these four BGCs reveals that, while microbes typically produce only one of the four, co-occurrence of *egt*/*sen* and *ovo*/*ovs* clusters are disproportionately high (**Fig. 3h**). In other words, there appears to be a selective advantage to producing both ergothioneine and selenoneine, or ovothiol and ovoselenol, as opposed to other combinations. This curious observation could point to the existence of ergothioneine/selenoneine and ovothiol/ovoselenol redox-couples, with the Se-containing compound providing the more reducing metabolite. Consistent with this idea is the one-electron peak oxidation potential of ovoselenol (**Fig. 3i**, **Supplementary Fig. 9**), which we measured by cyclic voltammetry (CV) to be ∼110 mV vs. the normal hydrogen electrode (NHE) and thus ∼340 mV lower than that of ovothiol^16^. As ovoselenol is significantly more reducing, it can act as a more effective antioxidant. The one-electron peak oxidation potentials of selenoneine (410 mV) and ergothioneine (810 mV) were also determined, with the latter similar to that of thiourea^17^. These trends are mirrored by the two-electron reduction potentials reported previously for these molecules: -490 mV^18^ and -60 mV^7^, respectively. While we could not determine the two-electron reduction potential of ovoselenol due to the complexity of the CV traces^19^, the observed peak potentials are consistent with a model in which one TSI can reduce another thereby forming a small-molecule redox chain. As noted above, many organisms appear to produce more than one TSI, with notable examples including *Limnobacter thiooxidans*, which encodes all four chemotypes (**Supplementary Fig. 3**). It is thus unlikely that these molecules are biologically redundant or interchangeable, but rather serve distinct purposes in the cell.

### Structural characterization of OvsA

The reaction catalyzed by OvsA points to an interesting convergence along the evolutionary trajectory of the NHISS family. Specifically, OvsA, despite sharing a recent common ancestor with the SenA/EgtB subfamilies, displays a regioselectivity similar to that of OvoA, an enzyme with which it shares only very little sequence similarity (**Supplementary Fig. 1**). We therefore wondered how the evolved architecture of the OvsA active site effects this convergence. The most striking differences between the OvsA subfamily’s composite motif and that of the other known subfamilies lie among the residues putatively involved in histidine/hercynine binding (**Fig. 2f**). For example, OvoA and OvsA, the two histidine-accepting members, feature a conserved alanine residue (A431 in OvsA) in place of the bulky aromatic residue common to hercynine-accepting subfamilies. To examine the function of these differences, we constructed an OvsA variant whose putative histidine-binding motif was altered to match that of SenA (OvsA**-**M401Y/Q430N/A431F, hereafter referred to as OvsA-YNF). These mutations introduced a strong preference for hercynine over histidine, as evidenced by competition assays in the presence of both histidine and hercynine (**Fig. 4a, b**). Moreover, selenylation of hercynine by OvsA-YNF was found to occur at the histidine 2′-carbon, suggesting that the *N*α-methylation pattern is linked to the substrate orientation within the enzymes’ active sites (**Fig. 4c**). Thus, three simple mutations can convert OvsA into SenA.

**Figure 4.**
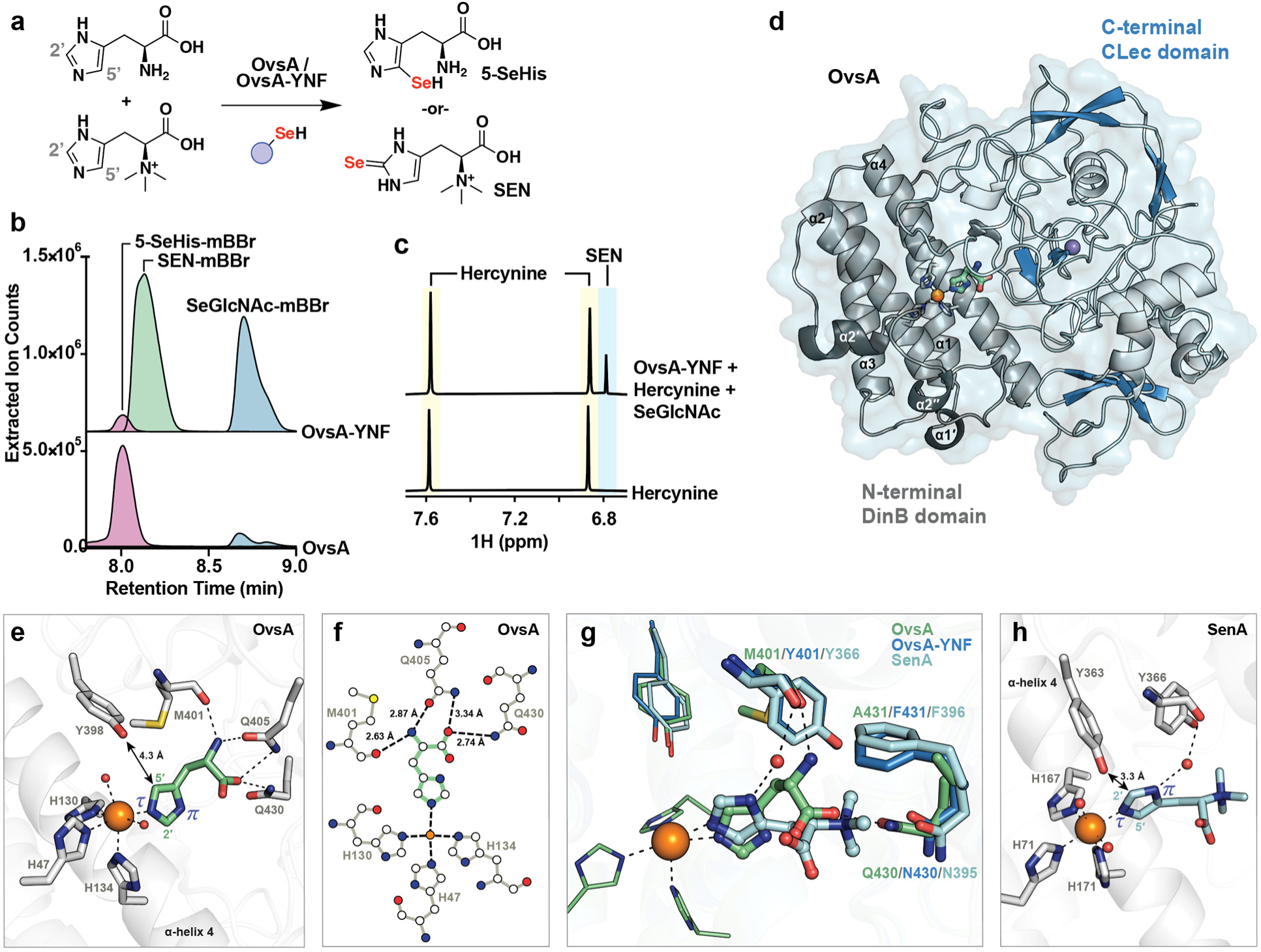
Mutagenesis and X-ray crystal structure analysis of OvsA reveal the basis for C–Se bond regioselectivity. (**a**) Competition assays demonstrate substrate preference and regioselectivity of OvsA and OvsA-M401Y/Q430N/A451F (OvsA-YNF, mutated to match the hercynine binding motif of SenA). A 1:1 mixture of histidine and hercynine is presented to either OvsA or OvsA-YNF in the presence of SeGlcNAc, and product compositions are determined. (**b**) Result of the competition assays in panel **a**. OvsA accepts histidine exclusively, while OvsA-YNF exhibits SenA-like activity, accepting mainly hercynine. (**c**) ^1^H-NMR analysis of the OvsA-YNF reaction with hercynine and SeGlcNAc, demonstrating 2′ C–Se bond formation to furnish selenoneine (SEN). (**d**) X-ray crystal structure of OvsA in complex with iron (orange) and histidine (green). (**e**) Active site architecture of OvsA reveals substrate interactions involved in the preference for 5′ C–Se bond formation. (**f**) 2D representation of OvsA substrate binding interactions. (**g**) Overlaid active sites of OvsA (green), OvsA-YNF (dark blue), and SenA (pale blue), illustrating the basis for imidazole reorientation. (**h**) Active site architecture of SenA in complex with iron and hercynine (PDB accession code 8K5J), illustrating imidazole reorientation for 2′ C–Se bond formation.

To acquire a more precise understanding of the OvsA active site architecture, we determined its X-ray crystal structure to 2.0 Å resolution (PDB accession code 8U42; **Supplementary Table 7**). The crystallographic model reveals that OvsA’s overall topology is very similar to other structurally characterized NHISS subfamilies, despite modest sequence identities of 25-35%. Alignment with its closest structural homolog, SenA, yields a Cα root-mean-square deviation (RMSD) of 1.8 Å (PDB accession code 8K5J; **Supplementary Table 8, Supplementary Fig. 10**). Broadly speaking, OvsA displays the representative NHISS fold, composed of an *N*-terminal DinB domain^20^ and a *C*-terminal domain with the formylglycine-generating enzyme (FGE) subclass of the C-type lectin (CLec) fold^21,22^ (see the Supporting Information for additional discussion of OvsA’s overall architecture). The catalytic iron center is coordinated by a three-His facial triad at the interface between these two domains. As in SenA, EgtB-I, and OvoA, the active site of OvsA additionally features a prominent Tyr residue (**Fig. 2e, f**). Previous mechanistic studies on EgtB-I and OvoA have demonstrated that this conserved residue is crucial for sulfoxide synthase activity^23-26^ and likely plays a similar role in the selenoxide synthases.

To further probe the structural basis for regioselectivity, we also determined the structure of OvsA complexed to histidine to 2.72 Å resolution (PDB accession code 8U41; **Fig. 4d, Supplementary Table 7, Supplementary Fig. 11**). Remarkably, the 5′-imidazole carbon is oriented toward the catalytic Tyr residue (Tyr398), thus facilitating C–Se bond formation at this position. The enzyme structure is preorganized to enable such reactivity and minimal changes to the overall scaffold are observed upon substrate binding, primarily in the loop-rich CLec domain and distant from the active site. This result is not entirely unexpected as SenA, EgtB-I, and EgtB-II demonstrate similarly fixed conformations even in the presence of substrate. Correct orientation of histidine in OvsA is enabled via four key hydrogen bonding interactions between the substrate’s amino and carboxylate groups and nearby residues: namely, the backbone carbonyl of Met401 (2.6 Å), as well as the side chains of Gln405 (2.9 and 3.3 Å) and Gln430 (2.7 Å) (**Fig. 4e, f**). The substrate is additionally coordinated to the iron cofactor through its imidazole τ-nitrogen. In the absence of O_2_ and a selenosugar, the remaining iron coordination sites are occupied by two water molecules. In contrast to histidine binding by OvsA, hercynine binding in SenA is primarily accomplished through electrostatic interactions with the substrate’s *N*α-methyl groups, including dipolar contacts with the Asn395 side chain and cation-π interactions with Phe396 (**Fig. 4h**). These features are recapitulated in OvsA-YNF (Asn430 and Phe431, respectively) (**Fig. 4h**), the crystal structure of which we determined to 3.06 Å resolution (PDB accession code 8UX5; **Supplementary Table 7**). The resultant model closely aligns with that of wild-type OvsA, with the only discernable differences located in the active site. In SenA, hercynine is further stabilized by a water-mediated hydrogen bond between the backbone carbonyl of Tyr366 (Tyr401 in OvsA-YNF) and the imidazole π-nitrogen (**Fig. 4g**). This interaction orients hercynine such that its 2′-imidazole carbon faces the catalytic Tyr398, thus enabling substitution at this carbon. OvsA-YNF likely provides an additional level of stability through Gln405, the side chain of which reorients to interact with this same water molecule (**Supplementary Fig. 12**). In the absence of the *N*α-methyl groups and associated dipolar interactions, the direct hydrogen bond between histidine’s primary amino group and Met401 in wild-type OvsA results in rotation of the substrate backbone toward the *N*-terminal end of α-helix 4. This reorganization of the active site forces rotation of the imidazole ring to maintain ideal bond geometry with iron, thereby enabling OvsA’s observed 5′ regioselectivity. We therefore propose that regioselectivity of C–S/Se bond formation in these enzymes is dictated by the orientation of the imidazole ring with respect to the catalytic Tyr residue, which is influenced by histidine’s *N*α-methylation pattern.

## Discussion

We report the discovery of the novel selenometabolite ovoselenol, which completes a quartet of redox-active thio/selenoimidazoles that are synthesized by diverse microbes. In addition to expanding the short list of selenium-containing small molecules, ovoselenol also highlights an important evolutionary convergence within the NHISS family. The phylogeny of the NHISS family can be crudely described as an early bifurcation event leading to two main branches, namely the OvoA-like enzymes (OvoA and EgtB-IV) and the EgtB-like enzymes (EgtB-I, EgtB-II, SenA, and OvsA). Despite the low sequence similarity between the extant members of the two branches (**Supplementary Fig. 1**), nature has found ways to repurpose certain subbranches such that they exhibit catalytic features of another. The first example of this phenomenon is EgtB-IV which, despite a remarkable similarity to OvoA, performs an EgtB-like reaction to furnish ergothioneine^11^. Herein, we identify the second such example in which an enzyme from the EgtB-like branch was evolutionarily remodeled, in this case, to generate the selenium isologue of ovothiol. The *ovs* cluster also encodes a standalone *N*π-methyltransferase reminiscent of the *C*-terminal domain of OvoA^8,14,27^ (**Supplementary Fig. 1**), suggesting another intriguing instance of gene exchange and repurposing among these biosynthetic pathways.

It is becoming increasingly evident that subtle differences among these enzymes are responsible for their distinct reactivities, which may not be obvious from sequence similarity alone. Important structural studies over the past decade have begun to illuminate the key residues responsible for engendering this catalytic diversity, affording us the ability to roughly predict the residues involved in substrate binding^10,13-15^. Informed by the structure of OvsA, we add an additional level of resolution to the prediction framework by compressing the sequences into composite motifs that can be used to classify and identify new families of NHISS enzymes. Indeed, there remain many family members with divergent motifs whose functions have yet to be uncovered (**Supplementary Fig. 2, Supplementary Table 4**).

Nature has clearly demonstrated a strong imperative toward generating all four chemotypes of histidine-derived thiols/selenols. Furthermore, in addition to repurposing of the NHISS scaffold, at least two other enzyme classes exist among anaerobic bacteria and archaea (EanB and MES) that assemble ergothioneine by completely different mechanisms^28,29^. The ubiquity of these pathways underscores the biological importance of this family of molecules, and the remarkable specificity of the enzymes involved implies potentially non-redundant biological functions for each of the final products. Finally, we hope that the addition of ovoselenol to the small-but-growing list of selenometabolites will inspire further studies to elucidate their roles in redox biology, and, more broadly catalyze further exploration of this emerging field.

## Methods

### Materials

All materials were purchased from Millipore-Sigma or Fisher Scientific unless otherwise specified. Cloning reagents were purchased from New England BioLabs (NEB). Codon-optimized gene fragments were purchased from Twist Biosciences. DNA primers were purchased from Sigma. Synthesis of N-acetyl-1-seleno-β-D-glucosamine (SeGlcNAc) was carried out as previously described^30,4^.

### General Experimental procedures

High-performance liquid chromatography-coupled mass spectrometry (HPLC-MS) was performed on an Agilent instrument equipped with a 1260 Infinity Series HPLC, an automated liquid sampler, a photodiode array detector, a JetStream ESI source, and the 6540 Series Q-tof mass spectrometer. HPLC-MS data were acquired and analyzed with Agilent MassHunter software. HPLC purifications were carried on an Agilent 1260 Infinity Series HPLC system equipped with a photodiode array detector and an automated fraction collector. Solvents for all LC-MS and HPLC experiments were water + 0.1% formic acid (Solvent A) and MeCN + 0.1% formic acid (Solvent B). HPLC data were acquired and analyzed with Agilent OpenLab software. Nuclear magnetic resonance (NMR) spectra were collected at the Princeton University Department of Chemistry’s NMR facility on a Bruker Avance III 500 MHz NMR spectrometer equipped with a DCH double resonance cryoprobe. NMR data were acquired using Bruker Topspin software and processed/analyzed using MestReNova software.

### Bioinformatics

All NHISS homologs were retrieved from the NCBI database by first performing initial blastp searches (E-value cutoff of 1e-30) using seed queries of previously characterized NHISS subfamilies (EgtB-I; EHI13262.1, EgtB-II; WP_014099805.1, SenA; WP_062361878.1, OvoA; CAO95049.1, EgtB-IV; WP_008197519.1). Next, 10 random, divergent sequences from each BLAST result were selected to be seeds for further blastp searches to capture the full diversity of NHISS homologs in the NCBI database. Following dereplication by strain name and removal of partial sequences, 39,678 total NHISS sequences remained.

Next, multiple sequence alignments (MSA) were performed separately on the hits for each subfamily query using MAFFT^31^ with either the L-INS-i algorithm (<1000 sequences) or the FFT-NS-2 algorithm (>1000 sequences). These MSAs were then used to extract the residue positions corresponding to iron-binding, histidine/hercynine-binding, thiol/selenol-binding, and catalytic tyrosine motifs as reported in previous crystal structure analyses of EgtB-I^13^, EgtB-II^10^, OvoA^14^, and SenA^15^. Residues at these 17 positions were then collapsed into a single composite motif for each retrieved NHISS. All sequences were then categorized into one of the following six subfamilies (or other) based on the following composite motif criteria, where “x” denotes any amino acid and “-” denotes a gap in the MSA:

- EgtB-I: xSDH--RRHQHYYNWDR (12,211 total sequences)
- EgtB-II: xSHHYYRRHQHxYNFFx (16,304 total sequences)
- SenA: xNEH--HRHxHYYN(F/Y)(F/Y)R (1,514 total sequences)
- OvsA: xNEH--HRHxHYMQAYR (228 total sequences)
- OvoA: RxYH--DxHxHYF(Y/F)AFx (4,375 total sequences)
- EgtB-IV: RxYH--DxHxHYxNWFx (466 total sequences)
- Other (4,580 total sequences)

Logo plots of composite motifs were generated using the Logomaker python script^32^.

### Phylogenetic tree construction

For the tree in **Fig. 2**, all full-length sequences were clustered at 70% sequence identity using the CD-hit clustering tool^33^, followed by random selection of ten representatives from each subfamily to be used for tree construction. For the tree in **Supplementary Fig. 2**, six of the most abundant uncharacterized subfamilies (grouped according to composite motifs) were selected for display, and five members of each of these subfamilies were used along with single members of the six characterized subfamilies for tree construction.

Phylogenetic trees were constructed by first generating MSAs using MAFFT with the L-INS-i algorithm, which were then used as input to FastTree^34^ for maximum-likelihood phylogeny approximation with parameters -gamma, -mlacc 2, and -slownii. Trees were visualized using FigTree (http://tree.bio.ed.ac.uk/software/figtree). The initial bifurcation between the OvoA-like (OvoA, EgtB-IV) and EgtB-like (EgtB-I, EgtB-II, SenA, OvsA) branches was selected as the tree root.

### Strains, media, and general culture conditions

Strains used in selenometabolite characterization and recombinant protein production experiments are listed in **Supplementary Table 1**. Bacto marine broth (Difco 2216) and agar plates were used for general maintenance and liquid cultures of *Marinobacter vinifirmus* DSM 17747. A growth medium consisting of 100 g/L NaCl, 5.4 g/L KCl, 10 g/L peptone, 2.5 g/L sodium acetate, 2 g/L yeast extract, 100 mM MgSO_4_, 100 mM MgCl_2_, 5 mM CaCl_2_ (+ 2 g/L agar for solid media), adjusted to pH 7.5 with Tris-HCl, was used for general maintenance and liquid cultures of *Halomonas utahensis* DSM 3051.

### Expression and purification of 6xHis-tagged OvsA, OvsM, and OvsA-YNF

Genes encoding OvsA, OvsM, and OvsA-M401Y/Q430N/A431F (OvsA-YNF) from *Halomonas utahensis* DSM 3051 were obtained as synthetic DNA fragments, codon-optimized for expression in *E. coli* and with overhangs to allow assembly into pET28b(+). Protein expression plasmids were assembled from gene fragments and vector pET28b(+), linearized with NdeI and XhoI (NEB), using HiFi DNA Assembly Master Mix (NEB) following the manufacturer’s instructions (**Supplementary Table 3**). Ligation mixtures were transformed into chemically-competent *E. coli* DH5α by heat-shock and plated onto LB agar containing 50 mg/L kanamycin. After confirmation by Sanger sequencing, assembled plasmids were transformed into *E. coli* BL21(DE3) for protein expression. For protein expression, starter cultures were prepared by inoculating 15 mL of LB medium containing 50 mg/L kanamycin with a single colony of *E. coli* BL21(DE3) carrying the desired plasmid. After overnight growth at 37 °C/200 rpm, starters were used to inoculate 2.8 L baffled Fernbach flasks containing 1.5 L Luria-Bertani (LB) broth supplemented with 50 mg/L kanamycin (1% inoculum) and incubated at 37 °C/200 rpm. At OD_600_ = 0.5–0.6, protein expression was induced with 0.2 mM IPTG, and cultures were incubated at 18 °C/200 rpm for an additional 12– 24 hours. Cells were pelleted by centrifugation (8,000 g, 15 min, 4 °C). The cell pastes were stored at -80 °C until purification.

All purification steps were carried out in a cold room at 4 °C. Cells were resuspended in lysis buffer (5 mL/g cell paste), which consisted of 25 mM Tris-HCl (pH 8), 500 mM NaCl, 10 mM imidazole, 10% glycerol, supplemented with 1 mM phenylmethylsulfonyl fluoride. Once homogenous, 0.1 mg/mL deoxyribonuclease I (Alfa Aesar) was added, and the cells were lysed by the addition of 5 mg/mL lysozyme followed by sonication using 30% power (∼150 W) in 15 s on/15 s off cycles for a total of 4 min. This process was repeated twice. The lysate was then clarified by centrifugation (17,000 g, 15 min, 4 °C) and loaded onto a 5 mL Ni-NTA column pre-equilibrated in lysis buffer. The column was washed with lysis buffer and His-tagged proteins were eluted with elution buffer consisting of 25 mM Tris-HCl (pH 8), 500 mM NaCl, 300 mM imidazole, 10% glycerol. Eluted proteins were then buffer-exchanged using a 50 mL column of Sephadex G-25 (Cytiva) into storage buffer consisting of 50 mM Tris-HCl (pH 8), 150 mM NaCl. For crystallography, OvsA and OvsA-YNF were further purified by fractionation on a Sephacryl S-200 HR HiPrep 16/60 column (Cytiva) with a running buffer consisting of 50 mM Tris-HCl (pH 8), 150 mM NaCl. Purified proteins were stored at -80 °C. Protein concentrations were determined spectrophotometrically on a Cary 60 UV-visible spectrophotometer (Agilent) using calculated molar extinction coefficients at 280 nm.

### Enzyme activity assays

Assays were carried out in 100 μL reactions containing 50 mM Tris-HCl (pH 8) and freshly prepared dithiothreitol (DTT) at a final concentration of 4 mM. Substrates were added at final concentrations of 1 mM. Enzymes were added at final concentrations of 10 μM. Reactions were initiated by the addition of SeGlcNAc (1 mM, monomer basis). Control assays were carried out in an identical fashion, except lacking one or more of the substrates/enzymes. Competition assays were performed similarly, except a mixture of 1 mM histidine and 1 mM hercynine was presented to either OvsA or OvsA-YNF.

After a 1-hour incubation period at room temperature, 50 µL of each reaction mixture was quenched with 50 µL of MeOH, while another 50 µL was derivatized with 50 µL of 10 mM monobromobimane (mBBr) in MeCN. Reactions were incubated for an additional 30 min at room temperature in the dark to allow for complete derivatization with mBBr. The samples were then filtered and analyzed by LC-MS using a Synergi Hydro-RP column (Phenomenex, 250 x 4.6 mm, 4 µm) with a flow rate of 1 mL/min. Elution programs for MeOH-quenched samples consisted of 5% solvent B for 4 min, followed by a gradient of 5–80% solvent B over 1 min, followed by a gradient of 80–100% over 1 min, and a hold at 100% for 4 min. Elution programs for mBBr-derivatized samples consisted of 5% solvent B for 3 min, followed by a gradient of 5–75% solvent B over 6 min, followed by a gradient of 75–100% over 1 min, and a hold at 100% for 4 min.

The assay to demonstrate 2′ C–Se bond formation by OvsA-YNF was carried out in 700 µL 50 mM Tris-HCl (pH 8) containing 4 mM DTT, 1 mM hercynine, 1 mM SeGlcNAc, and 10 μM OvsA. After 1 hour, protein was removed by centrifugal spin filter (10 kDa MWCO) and the sample was lyophilized, redissolved in 130 μL D_2_O, and analyzed by ^1^H NMR spectroscopy. Appearance of a single imidazole proton signal at 6.8 ppm indicates the formation of selenoneine, thus demonstrating the SenA-like regioselectivity of OvsA-YNF.

### Purification and structural characterization of 5-selenohistidine (5-SeHis) diselenide

A 70-mL enzymatic reaction containing 50 mM Tris-HCl (pH 8), 10 mM DTT, 2 mM histidine, 4 mM SeGlcNAc (monomer basis) and 10 μM OvsA was incubated at room temperature for 3 hours, followed by removal of protein by centrifugal spin filter (10 kDa MWCO). The crude reaction mixture was lyophilized, redissolved in water, and fractionated on a Luna Omega Polar C18 column (Phenomenex, 150 x 21.2 mm, 5 µm) with a flow rate of 15 mL/min and 100% solvent A. Fractions containing 5-SeHis were further purified by fractionation on an XBridge BEH Amide OBD column (Waters, 10 x 250 mm, 5 µm) with a flow rate of 4 mL/min and 60% solvent B. Purified 5-SeHis diselenide was dissolved in D_2_O and analyzed by NMR spectroscopy. The selenium atom was confirmed to be positioned at the imidazole C5 carbon, as evidenced by the absence of an imidazole C5 proton and diagnostic ^1^H-^13^C HMBC correlations. Full spectra, chemical shift assignments, and select 2D correlations can be found in **Supplementary Fig. 7** and **Supplementary Table 5**.

### Purification and structural characterization of ovoselenol (OVS) diselenide

A 70-mL enzymatic reaction containing 50 mM Tris-HCl (pH 8), 10 mM DTT, 2 mM histidine, 3 mM SAM, 4 mM SeGlcNAc (monomer basis), 10 μM OvsA, and 10 μM OvsM was incubated at room temperature for 3 hours, followed by removal of protein by centrifugal spin filter (10 kDa MWCO). The sample was acidified to pH 2 with concentrated HCl and loaded onto a Dowex 50WX8 cation exchange column preequilibrated with water. The column was washed with water until the elution pH was neutral, after which the column was eluted with 50 mL portions of 1%, 5%, 10%, 15%, and 20% NH_4_OH, sequentially. The 5% NH_4_OH fraction containing OVS diselenide was lyophilized, redissolved in water, and fractionated on a Luna Omega Polar C18 column (Phenomenex, 150 x 21.2 mm, 5 µm) with a flow rate of 15 mL/min and 100% solvent A. Fractions containing OVS diselenide were further purified by fractionation on an XBridge BEH Amide OBD column (Waters, 10 x 250 mm, 5 µm) with a flow rate of 4 mL/min and 60% solvent B. Purified OVS diselenide was dissolved in D_2_O and analyzed by NMR spectroscopy. Full spectra, chemical shift assignments, and select 2D correlations can be found in **Supplementary Fig. 8** and **Supplementary Table 6**.

### Disruption of the *ovsA* gene in *M. vinifirmus*

The *ovsA* gene in *M. vinifirmus* was disrupted by replacement of an internal region of the gene with a chloramphenicol resistant marker (Cm^R^) by homologous recombination. Genomic DNA from *Marinobacter vinifirmus* DSM 17747 was isolated using the Wizard Genomic DNA Purification Kit (Promega) following the manufacturer’s instructions. From genomic DNA, two ∼2kb regions flanking the *ovsA* gene were amplified using primer pairs Vini-Left-F/Vini-Left-R and Vini-Right-F/Vini-Right-R. The Cm^R^ gene was amplified from plasmid pGro7 (Takara Bio) using the primer pair Cm-Vini-F/Cm-Vini-R. Primers were designed such that amplicons contain overhangs to allow for assembly into pEX18Tet-SacB, a conjugative suicide vector encoding the *sacB* gene for counterselection^35^ (**Supplementary Table 2**). Flanking regions and Cm^R^ gene fragments were assembled into pEX18Tet-SacB linearized with HindIII and BamHI using HiFi DNA Assembly Master Mix. Ligations were transformed into chemically-competent *E. coli* DH5α by heat-shock and plated onto LB agar containing 25 mg/L Cm. After confirmation by Sanger sequencing, assembled plasmids were transformed into the conjugation donor *E. coli* JV36^36^.

For conjugation, single colonies of *M. vinifirmus* and *E. coli* JV36 containing the gene disruption vector were inoculated into 5 mL marine broth and 5 mL LB supplemented with 25 mg/L Cm, respectively, and placed at 37 °C/200 rpm. After 12 hours, both strains were pelleted by centrifugation and resuspended in 300 μL marine broth. A conjugation mixture containing 50 μL donor and 50 μL recipient was spotted onto marine agar and placed at 37 °C. After 24 hours, the spot was scraped from the plate, resuspended in 500 μL marine broth, and 1 μL of the mixture was plated onto marine agar supplemented with 25 mg/L Cm and 10% sucrose to select for double crossover mutants. Single colonies were restreaked once more on the same medium. The mutant genotype was verified by PCR with the primer pair Vini-KOcheck-F/ViniKOcheck-R (**Supplementary Table 2, Supplementary Fig. 6**).

### Selenometabolite production screens

For each strain tested, a single colony from an agar plate was inoculated into a sterile culture tube containing 5 mL of liquid medium and incubated at 37 °C/200 rpm. The starter cultures were then used to inoculate 25 mL liquid cultures supplemented with 50 μM of filter-sterilized Na_2_SeO_3_ and incubated at 37 °C/200 rpm. Production cultures of *H. utahensis* were grown for 3 days and *M. vinifirmus* (wild-type and Δ*ovsA*) for 16 hours. Following incubation, cultures were pelleted by centrifugation, resuspended in 300 μL MeCN containing 10 mM mBBr, and sonicated for 30 min at room temperature to facilitate cell lysis and selenol derivatization. Samples were pelleted by centrifugation and supernatants were analyzed by HPLC-MS. Analytes were separated on a Synergi Fusion-RP column (Phenomenex, 100 x 4.6 mm, 4 µm) with a flow rate of 0.5 mL/min and an elution program consisting of 5% solvent B for 3 min, followed by a gradient of 5–75% solvent B over 6 min, then a gradient of 75–100% over 1 min, and a hold at 100% for 4 min.

### Enzymatic preparation of selenoneine (SEN) diselenide

To obtain pure material for electrochemical measurements, SEN was purified from a large-scale reaction containing SenA and its substrates hercynine and SeGlcNAc. SenA from *Variovorax paradoxus* was expressed and purified as previously described^4^. A 60-mL enzymatic reaction containing 50 mM Tris-HCl (pH 8), 20 mM DTT, 5 mM hercynine, 6 mM SeGlcNAc (monomer basis), and 20 μM SenA was incubated at room temperature for 24 hours, followed by removal of protein by centrifugal spin filter (10 kDa MWCO). The sample was acidified to pH 2 with concentrated HCl and loaded onto a Dowex 50WX8 cation exchange column preequilibrated with water. The column was washed with water until the elution pH was neutral, after which the column was eluted with 50 mL portions of 1%, 5%, 10%, 15%, and 20% NH_4_OH, sequentially. The 5% NH_4_OH fraction containing SEN diselenide was lyophilized, redissolved in water, and purified on a Luna Omega Polar C18 column (Phenomenex, 150 x 21.2 mm, 5 µm) with a flow rate of 15 mL/min and 100% solvent A. NMR characterization: ^1^H NMR (500 MHz, D_2_O) δ 7.02 (s, 1H), 3.80 (dd, J = 11.7, 3.9 Hz, 1H), 3.11-3.24 (m, 2H), 3.17 (s, 9H); ^13^C NMR (126 MHz, D_2_O) δ 170.6, 134.9, 131.2, 120.8, 78.1, 52.1, 24.9.

### Peak oxidation potential measurements

Cyclic voltammetry (CV) measurements were performed with a CH Instruments 760C potentiostat running CHI600E software using a glassy carbon (3 mm diameter) working electrode, a Pt wire counter electrode, and an Ag wire pseudo-reference electrode in a conventional three-electrode cell. All cyclic voltammograms were collected in potassium phosphate buffer (0.1 M, pH 7.0). Solution containing the compound of interest was prepared at 2 mg/10 mL. The scan rate was 100 mV/s unless otherwise noted. All measurements were conducted at room temperature under an argon atmosphere. The glassy carbon working electrode was polished between measurements with an aluminum slurry on a microcloth polishing pad, followed by solvent rinses, and drying under a stream of nitrogen.

To reduce the diselenide form of ovoselenol and selenoneine, a solution containing either molecule was degassed under argon for 10 min and kept under a positive argon atmosphere. It was then subjected to programmed electrolysis (constant potential -0.55 V vs SCE, 30 min) using the same electrode assembly. Immediately following bulk electrolysis, linear sweep voltammetry data were collected. The potential of the pseudo-reference electrode was determined using the ferrocenium/ferrocene redox couple as an internal standard in anhydrous CH_3_CN, previously distilled and kept over molecular sieves and K_2_CO_3_. Tetrabutylammonium hexafluorophosphate (*n*Bu_4_NPF_6_, 0.1 M in CH_3_CN) was used as the supporting electrolyte. Measured potentials were adjusted to the saturated calomel electrode (SCE) scale (with E_1/2_ taken to be 0.40 V vs SCE in CH_3_CN), and then converted to normal hydrogen electrode (NHE).

### Crystallization of OvsA

Crystals of OvsA were grown using the sitting drop vapor diffusion method at room temperature. All crystals were obtained with 27 mg/mL OvsA in 50 mM Tris (pH 8), 150 mM NaCl. Crystals of substrate-free OvsA were found to diffract more strongly when grown anaerobically in the ferrous oxidation state. Therefore, to obtain crystals of substrate-free OvsA, 2 molar equivalents (eq.) of (NH_4_)_2_Fe(SO_4_)_2_ was added to the protein solution, which was then sonicated in an ultrasonic water bath for 15 min to remove O_2_ and left in an anaerobic chamber for an additional 4 hours to ensure anaerobicity. The anaerobic protein solution was reduced with 10 mM sodium dithionite and crystallized by mixing 1:1 with a precipitant solution of 0.1 M sodium acetate (pH 4.8), 3.5 M sodium formate. To obtain crystals of OvsA bound to histidine, 5 molar eq. of (NH_4_)_2_Fe(SO_4_)_2_ and 20 mM histidine were added to OvsA, which was then mixed 1:1 with a precipitant solution of 0.1 M sodium acetate (pH 4.8), 3.9 M sodium formate. To obtain crystals of OvsA M401Y/Q430N/A431F (YNF), 5 molar eq. of (NH_4_)_2_Fe(SO_4_)_2_ and 20 mM histidine were added to the protein solution, which was then mixed 1:1 with a precipitant solution of 0.1 M sodium acetate (pH 4.6), 3.5 M sodium formate, 10 mM sarcosine.

Clear hexagonal rods appeared within 1 week for all conditions and were fully formed within 2 weeks. During crystal harvesting, the crystals were looped and briefly transferred into cryoprotectant before flash freezing in liquid N_2_. For the substrate-free crystals, the cryoprotectant was comprised of the precipitant with 28.3% (v/v) ethylene glycol. For the histidine-containing WT OvsA crystals and OvsA-YNF crystals, the cryoprotectant was comprised of the precipitant with 25% (v/v) ethylene glycol and 20 mM histidine.

### X-ray data collection and processing

All crystals were maintained at 100 K to minimize X-ray-induced damage during data collection. Diffraction images for wild-type OvsA were indexed and integrated using iMosflm,^37^ while XDS^38^ was used for OvsA-YNF. The intensity data were then merged, scaled, and converted to structure-factor amplitudes using *POINTLESS*, *AIMLESS*, and *CTRUNCATE* within the CCP4 suite^39-41^. Model building was conducted in Coot^42^, the structures refined in Phenix^43^, and model quality assessed using Molprobity^44^. Data processing and refinement statistics can be found in **Supplementary Table 7**. Figures depicting the structures were generated with PyMOL, using chain A from all structures^45^. The 2-D interaction diagram for histidine binding was generated with LigPlot+^46^.

*OvsA in complex with iron + histidine*. Diffraction data for the crystals with iron and histidine were collected at beamline 21-ID-F of the Advanced Photon Source (APS) at Argonne National Laboratory using a Rayonix MX300 detector. Images were collected sequentially (Δφ = 1°) with an incident wavelength of 0.9787 Å. The structure of OvsA was solved via molecular replacement with *PHASER* using the AlphaFold2 predicted structure as the search model^47^, after removing all residues from the model with per-residue confidence scores (pLDDT) below 90%. The histidine-bound OvsA structure (PDB accession code 8U41) is in the hexagonal P 6_5_ space group, contains two molecules in the asymmetric unit, and was refined to 2.72 Å resolution.

*OvsA in complex with iron*. Diffraction data for the substrate-free crystals were collected at beamline 23-ID-B of the APS using an Eiger X 16M (Dectris) detector. Images were collected sequentially (Δφ = 0.2°) with an incident wavelength of 1.0332 Å. The structure was solved via molecular replacement using the histidine-bound structure as the search model in *PHASER*. The substrate-free OvsA structure (PDB accession code 8U42) is in the orthorhombic P 2_1_ 2_1_ 2_1_ space group, contains two molecules in the asymmetric unit, and was refined to 2.0 Å resolution.

*OvsA M401Y/Q430N/A431F (OvsA-YNF) in complex with iron.* Diffraction data for OvsA-YNF crystals were collected at beamline ID7B2 of the Cornell High Energy Synchrotron Source using an Eiger2 16M (Dectris) detector. Images were collected sequentially (Δφ = 0.25°) with an incident wavelength of 0.9686 Å. Crystals grown with hercynine diffracted poorly, so data were instead collected on crystals grown in the presence of histidine, which resulted in better crystal growth and higher resolution diffraction. Density was not observed, however, for bound histidine in the OvsA-YNF active site. The structure was solved via molecular replacement using the high-resolution substrate-free OvsA structure as the search model in *PHASER*. The structure (PDB accession code 8UX5) is in the hexagonal P 6_5_ space group, contains two molecules in the asymmetric unit, and was refined to 3.06 Å resolution.

### Reporting Summary

Further information on research design is available in the Nature Research Reporting Summary linked to this paper.

### Data Availability Statement

Protein crystal structure coordinates have been deposited with the Protein Data Bank (PDB) under accession numbers 8U42 (OvsA), 8U41 (Ovs+His), and 8UX5 (OvsA-YNF). Experimental data supporting the conclusions of this study are available within the article and its Supplementary information. Sequences were retrieved from the NCBI Conserved Domain Database (https://www.ncbi.nlm.nih.gov/cdd/) and the NCBI Non-redundant Protein Database (https://www.ncbi.nlm.nih.gov/protein/). Raw experimental data and complete bioinformatic datasets can be made available upon reasonable request.

## Acknowledgments

We thank the Eli Lilly-Edward C. Taylor Fellowship in Chemistry (to C.M.K.), the National Science Foundation (Graduate Research Fellowship Program No. 1937971 to K.A.I. and NSF CAREER Award No. 184786 to M.R.S.), and the National Institutes of Health (grant R35 GM147557 to K.M.D. and grants R01 GM140034 and R01 DA056358 to M.R.S.) for financial support. This research used resources of the Advanced Photon Source (APS) and the Center for High-Energy X-ray Sciences (CHEXS). APS is a U.S. Department of Energy (DOE) Office of Science User Facility operated for the DOE Office of Science by Argonne National Laboratory under Contract No. DE-AC02-06CH11357. Use of the LS-CAT Sector 21 was supported by the Michigan Economic Development Corporation and the Michigan Technology Tri-Corridor (grant 085P1000817). GM/CA@APS has been funded by the National Cancer Institute (ACB-12002) and the National Institute of General Medical Sciences (AGM-12006, P30GM138396). The Eiger 16M detector at GM/CA-XSD was funded by NIH grant S10 OD012289. CHEXS is supported by the NSF award DMR-1829070, and the MacCHESS resource is supported by NIGMS award 1-P30-GM124166-01A1 and NYSTAR.

## Author contributions

C.M.K. and M.R.S. conceived of the idea for the study. C.M.K. performed all bioinformatic and biochemistry experiments. K.A.I. performed all structural biology experiments. V.Y. synthesized SeGlcNAc and performed electrochemical measurements. C.M.K., K.A.I., V.Y., K.M.D., and M.R.S. analyzed data and prepared the manuscript.

## Additional information

**Supplementary information**

This report contains Supplementary Text, Supplementary Tables 1-8, and Supplementary Figs. 1-12.

**Correspondence and requests for materials** should be addressed to Mohammad R. Seyedsayamdost

## References

1. Walsh, C. T. The chemical biology of sulfur. The Royal Society of Chemistry. (2020).

2. Reich, H. J., & Hondal, R. J. Why Nature Chose Selenium. ACS Chem. Biol. 11, 821–841 (2016). doi: 10.1021/acschembio.6b00031.

3. Dunbar, K. L., Scharf, D. H., Litomska, A., & Hertweck, C. Enzymatic Carbon–Sulfur Bond Formation in Natural Product Biosynthesis. Chem. Rev. 117, 5521–5577 (2017). doi: 10.1021/acs.chemrev.6b00697.

4. Kayrouz, C. M., Huang, J., Hauser, N., & Seyedsayamdost, M. R. Biosynthesis of selenium-containing small molecules in diverse microorganisms. Nature 610, 199–204 (2022). doi: 10.1038/s41586-022-05174-2.

5. Cordell, G. A., & Lamahewage, S. N. S. Ergothioneine, Ovothiol A, and Selenoneine—Histidine-Derived, Biologically Significant, Trace Global Alkaloids. Molecules 27, 2673 (2022). doi: 10.3390/molecules27092673.

6. Gründemann, D., Harlfinger, S., Golz, S., Geerts, A., Lazar, A., Berkels, R., Jung, N., Rubbert, A., & Schömig, E. Discovery of the ergothioneine transporter. Proc. Natl. Acad. Sci. U.S.A. 102, 5256–5261 (2022). doi: 10.1073/pnas.0408624102.

7. Cheah, I. K., & Halliwell, B. Ergothioneine; antioxidant potential, physiological function and role in disease. Biochimica et Biophysica Acta (BBA) - Molecular Basis of Disease 1822, 784–793 (2012). doi: 10.1016/j.bbadis.2011.09.017.

8. Braunshausen, A., & Seebeck, F. P. Identification and Characterization of the First Ovothiol Biosynthetic Enzyme. J. Am. Chem. Soc. 133, 1757–1759 (2011). doi: 10.1021/ja109378e.

9. Seebeck, F. P. In Vitro Reconstitution of Mycobacterial Ergothioneine Biosynthesis. J. Am. Chem. Soc. 132, 6632–6633 (2010). doi: 10.1021/ja101721e.

10. Stampfli, A. R., Goncharenko, K. V., Meury, M., Dubey, B. N., Schirmer, T., & Seebeck, F. P. An Alternative Active Site Architecture for O2 Activation in the Ergothioneine Biosynthetic EgtB from *Chloracidobacterium thermophilum*. J. Am. Chem. Soc. 141, 5275–5285 (2019). doi: 10.1021/jacs.8b13023.

11. Liao, C., & Seebeck, F. P. Convergent Evolution of Ergothioneine Biosynthesis in Cyanobacteria. ChemBioChem 18, 2115–2118 (2017). doi: 10.1002/cbic.201700354.

12. Hu, W., Song, H., Sae Her, A., Bak, D. W., Naowarojna, N., Elliott, S. J.,Qin, L, Chen, X., Liu, P. Bioinformatic and Biochemical Characterizations of C–S Bond Formation and Cleavage Enzymes in the Fungus Neurospora crassa Ergothioneine Biosynthetic Pathway. Org. Lett. 16, 5382–5385 (2014). doi: 10.1021/ol502596z.

13. Goncharenko, K. V., Vit, A., Blankenfeldt, W., & Seebeck, F. P. Structure of the Sulfoxide Synthase EgtB from the Ergothioneine Biosynthetic Pathway. Angew. Chem. Int. Ed. 54, 2821–2824 (2015). doi: 10.1002/anie.201410045.

14. Wang, X., Hu, S., Wang, J., Zhang, T., Ye, K., Wen, A., Zhu, G., Vegas, A., Zhang, L., Yan, W., Liu, X., & Liu, P. Biochemical and Structural Characterization of OvoATh2: A Mononuclear Nonheme Iron Enzyme from *Hydrogenimonas thermophila* for Ovothiol Biosynthesis. ACS Catal. 13, 15417–15426 (2023). doi: 10.1021/acscatal.3c04026.

15. Liu, M., Yang, Y., Huang, J.-W., Dai, L., Zheng, Y., Cheng, S., He, H., Chen, C.-C., Guo, R.-T. Structural insights into a novel nonheme iron-dependent oxygenase in selenoneine biosynthesis. Int. J. Biol. Macromol. 256, 128428 (2024). doi: 10.1016/j.ijbiomac.2023.128428.

16. Marjanovic, B., Simic, M. G. & Jovanovic, S. V. Heterocyclic thiols as antioxidants: why ovothiol C is a better antioxidant than ergothioneine. Free Radic. Biol. Med. 18, 679–685 (1995). doi: 10.1016/0891-5849(94)00186-n.

17. Kirchnerova, J., & Purdy, W. C. The mechanism of the electrochemical oxidation of thiourea. Anal. Chim. Acta 123, 83–95 (1981). doi: 10.1016/S0003-2670(01)83161-5

18. Yamashita, M., & Yamashita, Y. Selenoneine in Marine Organisms. Springer Berlin Heidelberg. 1059-1069 (2015).

19. Zhu, Q., Costentin, C., Stubbe, J., & Nocera DG. Disulfide radical anion as a super-reductant in biology and photoredox chemistry. Chem. Sci. 14, 6876–6881 (2023). doi: 10.1039/d3sc01867a

20. Cooper, D. R., Grelewska, K., Kim, C.-Y., Joachimiak, A., & Derewenda, Z. S. The structure of DinB from Geobacillus stearothermophilus: a representative of a unique four-helix-bundle superfamily. *Acta Crystallogr.*, Sect. F: Struct. Biol. Cryst. Commun. 66, 219–224 (2010). doi: 10.1107/S1744309109053913.

21. McMahon, S. A., Miller, J. L., Lawton, J. A., Kerkow, D. E., Hodes, A., Marti-Renom, M. A., Doulatov, S., Narayanan, E., Sali, A., Miller, J. F., & Ghosh, P. The C-type lectin fold as an evolutionary solution for massive sequence variation. Nat. Struct. Mol. Biol. 12, 886–892 (2005). doi: 10.1038/nsmb992.

22. Le Coq, J., & Ghosh, P. Conservation of the C-type lectin fold for massive sequence variation in a Treponema diversity-generating retroelement. Proc. Natl. Acad. Sci. U.S.A. 108, 14649–14653 (2011). doi: 10.1073/pnas.1105613108.

23. Chen, L., Naowarojna, N., Song, H., Wang, S., Wang, J., Deng, Z., Zhao, C., & Liu, P. Use of a Tyrosine Analogue To Modulate the Two Activities of a Nonheme Iron Enzyme OvoA in Ovothiol Biosynthesis, Cysteine Oxidation versus Oxidative C–S Bond Formation. J. Am. Chem. Soc. 140, 4604–4612 (2018). doi: 10.1021/jacs.7b13628.

24. Goncharenko, K. V., & Seebeck, F. P. Conversion of a non-heme iron-dependent sulfoxide synthase into a thiol dioxygenase by a single point mutation. Chem. Commun. 52, 1945–1948 (2016). doi: 10.1039/C5CC07772A.

25. Cheng, R., Weitz, A.C., Paris, J., Tang, Y., Zhang, J., Song, H., Naowarojna, N., Li, K., Qiao, L., Lopez, J., Grinstaff, M. W., Zhang, L., Guo, Y., Elliot, S, & Liu, P. OvoAMtht from Methyloversatilis thermotolerans ovothiol biosynthesis is a bifunction enzyme: thiol oxygenase and sulfoxide synthase activities. Chem. Sci. 13, 3589–3598 (2022). doi: 10.1039/D1SC05479A.

26. Chen, L., Naowarojna, N., Chen, B., Xu, M., Quill, M., Wang, J., Deng, Z., Zhao, C., & Liu, P. Mechanistic Studies of a Nonheme Iron Enzyme OvoA in Ovothiol Biosynthesis Using a Tyrosine Analogue, 2-Amino-3-(4-hydroxy-3-(methoxyl) phenyl) Propanoic Acid (MeOTyr). ACS Catal. 9, 253–258 (2019). doi: 10.1021/acscatal.8b03903.

27. Naowarojna, N., Huang, P., Cai, Y., Song, H., Wu, L., Cheng, R., Li, Y., Wang, S., Lyu, H., Zhang, L., Zhou, J., & Liu, P. In Vitro Reconstitution of the Remaining Steps in Ovothiol A Biosynthesis: C–S Lyase and Methyltransferase Reactions. Org. Lett. 20, 5427–5430 (2018). doi: 10.1021/acs.orglett.8b02332.

28. Burn, R., Misson, L., Meury, M., & Seebeck, F. P. Anaerobic Origin of Ergothioneine. Angew. Chem. 129, 12682–12685 (2017). doi: 10.1002/ange.201705932.

29. Beliaeva, M. A., & Seebeck, F. P. Discovery and Characterization of the Metallopterin-Dependent Ergothioneine Synthase from Caldithrix abyssi. JACS Au 2, 2098–2107 (2022). doi: 10.1021/jacsau.2c00365.

## Methods References

30. Kumar, A. A., Illyes, T. Z., Kover, K. E., & Szilagyi, L. Convenient syntheses of 1,2-trans selenoglycosides using isoselenuronium salts as glycosylselenenyl transfer reagents. Carbohydrate Res. 360, 8-18 (2012). doi: 10.1016/j.carres.2012.07.012.

31. Katoh, K. Mafft: A novel method for rapid multiple sequence alignment based on fast fourier transform. Nucleic Acids Res. 30, 3059–3066 (2002). doi: 10.1093/nar/gkf436.

32. Tareen, A., & Kinney, J. B. Logomaker: Beautiful sequence logos in python. Bioinformatics 36, 2272– 2274 (2019). doi: 10.1093/bioinformatics/btz921.

33. Fu, L., Niu, B., Zhu, Z., Wu, S., & Li, W. CD-hit: Accelerated for clustering the next-generation sequencing data. Bioinformatics 28, 3150–3152 (2012). doi: 10.1093/bioinformatics/bts565.

34. Price, M. N., Dehal, P. S., & Arkin, A. P. FastTree 2 – Approximately Maximum-Likelihood Trees for Large Alignments. PLoS One 5, e9490 (2010). doi: 10.1371/journal.pone.0009490.

35. Hoang, T. T., Karkhoff-Schweizer, R. R., Kutchma, A. J., & Schweizer, H. P. A broad-host-range Flp-FRT recombination system for site-specific excision of chromosomally-located DNA sequences: application for isolation of unmarked Pseudomonas aeruginosa mutants. Gene 212, 77–86 (1998). doi: 10.1016/S0378-1119(98)00130-9.

36. Blodgett, J. A. V., Oh, D.-C., Cao, S., Currie, C. R., Kolter, R., & Clardy, J. Common biosynthetic origins for polycyclic tetramate macrolactams from phylogenetically diverse bacteria. Proc. Natl. Acad. Sci. U.S.A. 107, 11692–11697 (2010). doi: 10.1073/pnas.1001513107.

37. Battye, T. G. G., Kontogiannis, L., Johnson, O., Powell, H. R., & Leslie, A. G. W. iMOSFLM: a new graphical interface for diffraction-image processing with MOSFLM. *Acta Crystallogr.*, Sect. D: Biol. Crystallogr. 67, 271–281 (2011). doi: 10.1107/S0907444910048675.

38. Kabsch, W. XDS. Acta Crystallogr., Sect. D: Biol. Crystallogr. 66, 125 (2010). doi: 10.1107/S0907444909047337.

39. Evans, P. R. An introduction to data reduction: space-group determination, scaling and intensity statistics. *Acta Crystallogr.*, Sect. D: Biol. Crystallogr. 67, 282–292 (2011). doi: 10.1107/S090744491003982X.

40. Evans, P. R., & Murshudov, G. N. How good are my data and what is the resolution? *Acta Crystallogr.*, Sect. D: Biol. Crystallogr. 69, 1204–1214 (2013). doi: 10.1107/S0907444913000061.

41. Winn, M.D., Ballard, C.C., Cowtan, K.D., Dodson, E.J., Emsley, P., Evans, P.R., Keegan, R.M., Krissinel, E.B., Leslie, A.G., McCoy, A., et al. Overview of the CCP4 suite and current developments. Acta Crystallogr. D Biol. Crystallogr. 67, 235-242 (2012). doi: 10.1107/S0907444910045749.

42. Emsley, P., Lohkamp, B., Scott, W. G., & Cowtan, K. Features and development of Coot. Acta Crystallogr., Sect. D: Biol. Crystallogr. 66, 486–501 (2010). doi: 10.1107/S0907444910007493.

43. Adams, P.D., Afonine, P.V., Bunkoczi, G., Chen, V.B., Davis, I.W., Echols, N., Headd, J.J., Hung, L.W., Kapral, G.J., Grosse-Kunstleve, R.W., et al. PHENIX: a comprehensive Python-based system for macromolecular structure solution. Acta Crystallogr., Sect. D: Biol. Crystallogr. 66, 213–221 (2010). doi: 10.1107/S0907444909052925.

44. Williams, C.J., Headd, J.J., Moriarty, N.W., Prisant, M.G., Videau, L.L., Deis, L.N., Verma, V., Keedy, D.A., Hintze, B.J., Chen, V.B., et al. MolProbity: More and better reference data for improved all-atom structure validation. Protein Sci. 27, 293–315 (2018). doi: 10.1002/pro.3330.

45. Schrodinger, LLC (2023). The PyMOL molecular graphics system, version 2.5.5.

46. Laskowski, R. A., & Swindells, M. B. LigPlot+: Multiple Ligand–Protein Interaction Diagrams for Drug Discovery. J. Chem. Inf. Model. 51, 2778–2786 (2011). doi: 10.1021/ci200227u.

47. Jumper, J., Evans, R., Pritzel, A., Green, T., Figurnov, M., Ronneberger, O., Tunyasuvunakool, K., Bates, R., Zidek, A., Potapenko, A., et al. Highly accurate protein structure prediction with AlphaFold. Nature 596, 583–589 (2021). doi: 10.1038/s41586-021-03819-2.

