## Supporting Information for "Ovoselenol, a Selenium-containing Antioxidant Derived from Convergent Evolution"

**Table of Contents**


| | |
|---|---|
| Coding DNA sequences and amino acid sequences of recombinant proteins | Pages S3-S4 |
| Supplementary Table 1 | Page S5 |
| Supplementary Table 2 | Page S6 |
| Supplementary Table 3 | Page S6 |
| Supplementary Table 4 | Page S7 |
| Supplementary Table 5 | Page S8 |
| Supplementary Table 6 | Page S8 |
| Supplementary Table 7 | Page S9 |
| Supplementary Table 8 | Page S10 |
| Supplementary Figure 1 | Page S11 |
| Supplementary Figure 2 | Page S12 |
| Supplementary Figure 3 | Page S13 |
| Supplementary Figure 4 | Page S14 |
| Supplementary Figure 5 | Page S15 |
| Supplementary Figure 6 | Page S16 |
| Supplementary Figure 7 | Pages S17-S20 |
| Supplementary Figure 8 | Pages S21-S23 |
| Additional Discussion of OvsA Structure | Page S24 |
| Supplementary Figure 9 | Page S25 |
| Supplementary Figure 10 | Page S26 |
| Supplementary Figure 11 | Page S26 |
| Supplementary Figure 12 | Page S27 |
| Supplementary References | Page S28 |

**Coding DNA sequences and amino acid sequences of recombinant proteins**

**6xHis-*ovsA* CDS (from *H. utahensis*, synthesized, codon-optimized)**
ATGGGCAGCAGCCATCATCATCATCATCACAGCAGCGGCCTGGTGCCGCGCGGCAGCCATATGAACGAC
CGCGAAAGTCTGATCCAGGCACTTCATCATACACGTGATCGGGTGAAAGATCTGGTTTGTTCATTGCGTG
AAGACCAGTTAAGTGTACCGTATCATCCGGGTGTTAACCCTCCGGTTTGGGAAATGGGCCATTCAACCTT
CTTCTATGAGGTTTTTGTTCTTAACTGGCTTGACGGCACGCCGAGCTATGATCCTTCTATGGATGACTTGT
GGGACAGCTTTCATATGGATCATGAAGATAGATGGTCTAAAACTCTGTTCCCCTCACGCGAAGATACCCT
TGCGTATATGGACACGATTATTCAGCGTATGGAGGATCGAATTCGCAACCAGCCACTCACGGATGAAGC
ACTCTACTTATACCGTTATGCCATCTATCACCAAAATATGCATGTTGAATCAATGACTTGGTGTCGTCAAA
CTGTTGGATACCCTGCCCCCCCGTTCGCTGAGCCGAAAGGTCTGACTGGTGAGGGTGTTGATCAGGATG
CGCGGGGTGACGCAACCATTCCGGCCGGTCGCTATCTTATCGGCCTGCCAGCAAACCGTGATAGCGATG
CTTACGCGACCGAAGATTTCGGCTTTGATAACGAAAAACCGGCCTTCGAAGTGGATATGCCTGAATTCTC
TATCTCACGGACACTCGTAACAAACGGCGAATTTCAGAAATTCGTTGAAGAAGGAGGCTATGAGAGACC
CGAGTTCTGGTCACAGGGAGGCAGAAAATGGCTGGAGCGCGAAATCAATCTGAATTTTGGCTCCGGTG
AGCCACCACTTATGGGGCGTCAAACCCATCCGTTCCATTGGCGGAAGCGCGATGGACGTTGGTATGAGC
GAGTCTTTGACCAGTGGTTACCGCTGGAACCCGGTCACCCTGTCAAGCAGATTTCGTATTGGGAAGCTG
AGGCATTTTGCGCTTGGGCGGGACGGCGCTTGCCATCTGAATATGAGTGGGAAGTTGCCGCGCTTGCAA
ATAAGCCTGGTGAAGAACGTCGCCGTTATCCATGGGGTAACGAGATGGATCCTGCGAAGTTAGATATG
GATCAGCGATATATGGGCCGCGTCCCTGTCACAGCTTTTCCGGCCGGTGAGTCACCATTTGGCTGTCGCC
AAATGCTTGGTACCGTATGGGAGTGGACGGGAAACCAGTTTATGCCTTATGACGGATTTTCAGTTGACA
TGTATCCTTTCATGTCGACTCTGCAGTTTGCCACGCATAAAACCACTAAAGGCGGAGGCTGTGCGGCATC
CTCTATGCTGATTCGGGGTACTTATCGTCAAGCTTACCACCCTGACCGATGCGATGTATACACCGGTTTC
AGAACATGCGCCCTGTCATCTCAGGATTAA

**6xHis-OvsA protein sequence**
MGSSHHHHHHSSGLVPRGSHMNDRESLIQALHHTRDRVKDLVCSLREDQLSVPYHPGVNPPVWEMGHST
FFYEVFVLNWLDGTPSYDPSMDDLWDSFHMDHEDRWSKTLFPSREDTLAYMDTIIQRMEDRIRNQPLTDE
ALYLYRYAIYHQNMHVESMTWCRQTVGYPAPPFAEPKGLTGEGVDQDARGDATIPAGRYLIGLPANRDSDA
YATEDFGFDNEKPAFEVDMPEFSISRTLVTNGEFQKFVEEGGYERPEFWSQGGRKWLEREINLNFGSGEPPL
MGRQTHPFHWRKRDGRWYERVFDQWLPLEPGHPVKQISYWEAEAFCAWAGRRLPSEYEWEVAALANKP
GEERRRYPWGNEMDPAKLDMDQRYMGRVPVTAFPAGESPFGCRQMLGTVWEWTGNQFMPYDGFSVD
MYPFMSTLQFATHKTTKGGGCAASSMLIRGTYRQAYHPDRCDVYTGFRTCALSSQD*

**6xHis-*ovsM* CDS (from *H. utahensis*, synthesized, codon-optimized)**
ATGGGCAGCAGCCATCATCATCATCATCACAGCAGCGGCCTGGTGCCGCGCGGCAGCCATATGACCGAT
TTTTATGAAAGCGATGAACAGCTGGGTCAGTATATGGATTTTCATTATGGCCCGGAACATTTTGGCGTGC
CGAACTTTCCCAAAACCTGCGCGGAACGCTGCCTGCAAGCGCGTCCGCGTGGCAATCGGGCGCTGGATC
TGGGTTGTGCGACCGGCCGAAGCACCCTGGAACTGGCGCGCGGCTTTGGCCATGTGCTGGGCATGGAT
CTGAGCCATCGCTTTATTGATGCGGCGGAACGCCTGCGCCGCGATGGCGAACTGACCTATAGCCTGGTG
GATGAAGGCGAAATTGGCCATCAAGAAACCGCGAGCCTGGCGGAACTGGGCCTGGCGGGCGAAGCGG
CGCGCGTGAGCTTTGAAGAATGCGATGCTGGCAACCTGGGCCCGGAATATAGCGGCTATGATCTGATTT
TTGCCGGCAACCTGATTGACCGTATGCCGAACCCGGGCCCGTTTCTCAGCGGCTTGAGGGAACGTCTGA
ATCCGGGCGGCGTGCTGGTGGTGACGAGCCCGTATACCCTGCTGCCGGAATTTACCCCGCGCGAAAGCT

GGATTGGCGGCTATTATGATGCGGATGGCAAACCGGTGACCGTGCTGGATGGCATGCGCGCGCATCTG
GAACCGGGCATGCAGCTGGCGGCGGAACCGGAAGATGTGCCGTTTGTGATTCGCGAAACCCGCCGCAA
ACATCAGCATACCCGCGCGCAGCTGACCGTGTGGGAACGCAAAGATTAA

**6xHis-OvsM protein sequence**
MGSSHHHHHHSSGLVPRGSHMTDFYESDEQLGQYMDFHYGPEHFGVPNFPKTCAERCLQARPRGNRALD
LGCATGRSTLELARGFGHVLGMDLSHRFIDAAERLRRDGELTYSLVDEGEIGHQETASLAELGLAGEAARVSF
EECDAGNLGPEYSGYDLIFAGNLIDRMPNPGPFLSGLRERLNPGGVLVVTSPYTLLPEFTPRESWIGGYYDAD
GKPVTVLDGMRAHLEPGMQLAAEPEDVPFVIRETRRKHQHTRAQLTVWERKD*

**6xHis-*ovsA*-YNF CDS (from *H. utahensis*, mutagenized, synthesized, codon-optimized)**
ATGGGCAGCAGCCATCATCATCATCATCACAGCAGCGGCCTGGTGCCGCGCGGCAGCCATATGAACGAC
CGCGAAAGTCTGATCCAGGCACTTCATCATACACGTGATCGGGTGAAAGATCTGGTTTGTTCATTGCGTG
AAGACCAGTTAAGTGTACCGTATCATCCGGGTGTTAACCCTCCGGTTTGGGAAATGGGCCATTCAACCTT
CTTCTATGAGGTTTTTGTTCTTAACTGGCTTGACGGCACGCCGAGCTATGATCCTTCTATGGATGACTTGT
GGGACAGCTTTCATATGGATCATGAAGATAGATGGTCTAAAACTCTGTTCCCCTCACGCGAAGATACCCT
TGCGTATATGGACACGATTATTCAGCGTATGGAGGATCGAATTCGCAACCAGCCACTCACGGATGAAGC
ACTCTACTTATACCGTTATGCCATCTATCACCAAAATATGCATGTTGAATCAATGACTTGGTGTCGTCAAA
CTGTTGGATACCCTGCCCCCCCGTTCGCTGAGCCGAAAGGTCTGACTGGTGAGGGTGTTGATCAGGATG
CGCGGGGTGACGCAACCATTCCGGCCGGTCGCTATCTTATCGGCCTGCCAGCAAACCGTGATAGCGATG
CTTACGCGACCGAAGATTTCGGCTTTGATAACGAAAAACCGGCCTTCGAAGTGGATATGCCTGAATTCTC
TATCTCACGGACACTCGTAACAAACGGCGAATTTCAGAAATTCGTTGAAGAAGGAGGCTATGAGAGACC
CGAGTTCTGGTCACAGGGAGGCAGAAAATGGCTGGAGCGCGAAATCAATCTGAATTTTGGCTCCGGTG
AGCCACCACTTATGGGGCGTCAAACCCATCCGTTCCATTGGCGGAAGCGCGATGGACGTTGGTATGAGC
GAGTCTTTGACCAGTGGTTACCGCTGGAACCCGGTCACCCTGTCAAGCAGATTTCGTATTGGGAAGCTG
AGGCATTTTGCGCTTGGGCGGGACGGCGCTTGCCATCTGAATATGAGTGGGAAGTTGCCGCGCTTGCAA
ATAAGCCTGGTGAAGAACGTCGCCGTTATCCATGGGGTAACGAGATGGATCCTGCGAAGTTAGATATG
GATCAGCGATATATGGGCCGCGTCCCTGTCACAGCTTTTCCGGCCGGTGAGTCACCATTTGGCTGTCGCC
AAATGCTTGGTACCGTATGGGAGTGGACGGGAAACCAGTTTATGCCTTATGACGGATTTTCAGTTGACA
TGTATCCTTTCTATTCGACTCTGCAGTTTGCCACGCATAAAACCACTAAAGGCGGAGGCTGTGCGGCATC
CTCTATGCTGATTCGGGGTACTTATCGTAACTTCTACCACCCTGACCGATGCGATGTATACACCGGTTTCA
GAACATGCGCCCTGTCATCTCAGGATTAA

**6xHis-OvsA-YNF protein sequence**
MGSSHHHHHHSSGLVPRGSHMNDRESLIQALHHTRDRVKDLVCSLREDQLSVPYHPGVNPPVWEMGHST
FFYEVFVLNWLDGTPSYDPSMDDLWDSFHMDHEDRWSKTLFPSREDTLAYMDTIIQRMEDRIRNQPLTDE
ALYLYRYAIYHQNMHVESMTWCRQTVGYPAPPFAEPKGLTGEGVDQDARGDATIPAGRYLIGLPANRDSDA
YATEDFGFDNEKPAFEVDMPEFSISRTLVTNGEFQKFVEEGGYERPEFWSQGGRKWLEREINLNFGSGEPPL
MGRQTHPFHWRKRDGRWYERVFDQWLPLEPGHPVKQISYWEAEAFCAWAGRRLPSEYEWEVAALANKP
GEERRRYPWGNEMDPAKLDMDQRYMGRVPVTAFPAGESPFGCRQMLGTVWEWTGNQFMPYDGFSVD
MYPFYSTLQFATHKTTKGGGCAASSMLIRGTYRNFYHPDRCDVYTGFRTCALSSQD*

**Supplementary Table 1.** Bacterial strains used or generated in this study.

| **Strain** | **Purpose** | **Source** |
|---|---|---|
| *Halomonas utahensis* DSM 3051[a] | Gammaproteobacteria containing *ovs* cluster, production screening for OVS[a] | DSMZ[a] |
| *Marinobacter vinifirmus* DSM 17747 | Gammaproteobacteria containing *ovs* cluster, production screening for OVS | DSMZ |
| *Marinobacter vinifirmus* Δ*ovsA* | Disruption of *ovsA* gene for production screening of *ovs* pathway intermediates | This study |
| *E. coli* DH5α | Host strain for cloning | NEB |
| *E. coli* BL21(DE3) | Host strain for protein expression | NEB |
| *E. coli* BL21(DE3) + pET-28b(+)_6xHis-OvsA | Host strain for expression of 6xHis-OvsA | This study |
| *E. coli* BL21(DE3) + pET-28b(+)_6xHis-OvsM | Host strain for expression of 6xHis-OvsM | This study |
| *E. coli* BL21(DE3) + pET-28b(+)_6xHis-OvsA-M401Y/Q430N/A431F | Host strain for expression of 6xHis-OvsA-YNF | This study |
| *E. coli* JV36 | Conjugation donor for *M. vinifirmus* gene disruption | Ref[36] |

[a]*Halomonas utahensis* DSM 3051 is the strain name associated with the NCBI record of the genome containing *ovsA* (genome: NZ_JACHUZ010000000, OvsA: WP_077530365.1). However, the taxonomic affiliation of this strain was brought into question in 2006 and was swapped with *Halovibrio variabilis* DSM 3050 in 2014[48,49]. Therefore, the genome record of *H. utahensis* DSM 3051 in the NCBI database actually refers to the genome of the strain deposited as *H. variabilis* DSM 3050 in the DSMZ strain repository. For simplicity, we herein refer to the name associated with the NCBI record (*H. utahensis* DSM 3051). However, OVS production screening experiments were performed using the strain denoted *H. variabilis* DSM 3050 from the DSMZ collection.

**Supplementary Table 2.** Primers used in this work.

| Primer | Sequence (5′->3′) | Purpose |
|---|---|---|
| Vini-Left-F | TAAAACGACGGCCAGTGCCAGTGGTGGACTGGCTGTT AAG | Amplification of 5′ flanking arm for *ovsA* knockout in *M. vinifrmus* |
| Vini-Left-R | AGTGGCCCATTTCCCAGAG | |
| Vini-Right-F | AACCTGGTTCGTGGCACTTA | Amplification of 3′ flanking arm for *ovsA* knockout in *M. vinifrmus* |
| Vini-Right-R | ATTCGAGCTCGGTACCCGGGATGCCATCACCAACATC CAG | |
| Cm-Vini-F | GCTCTGGGAAATGGGCCACTTGATCGGCACGTAAGA GGTT | Amplification of $Cm^R$ from pGro7 for *ovsA* knockout in *M. vinifrmus* |
| Cm-Vini-R | TAAGTGCCACGAACCAGGTTTTACGCCCCGCCCTG | |
| Vini-KOcheck-F | ATCGAGGATCTGGCAGACTG | Verification of *ovsA* knockout in *M. vinifirmus* |
| Vini-KOcheck-R | TCCCATATCACCAGCTCACC | |

**Supplementary Table 3.** Plasmids used or generated in this study.

| Plasmid | Purpose | Source |
|---|---|---|
| pET-28b(+) | $Kan^R$; Expression vector | Novagen |
| pET-28b(+)_6xHis-OvsA | Expression of 6xHis-OvsA | This study |
| pET-28b(+)_6xHis-OvsM | Expression of 6xHis-OvsM | This study |
| pET-28b(+)_6xHis-OvsA-M401Y/Q430N/A431F | Expression of 6xHis-OvsA-YNF | This study |
| pEX18Tet-SacB | $Tet^R$, SacB; Conjugation vector for gene disruption | Ref[35] |
| pEX18Tet-SacB-ViniKO | $Tet^R$, SacB, $Cm^R$; Disruption of *ovsA* gene in *M. vinifirmus* | This study |
| pGro7 | Source of $Cm^R$ gene | Takara Bio |

**Supplementary Table 4.** Additional information about characterized and select uncharacterized NHISS subfamilies. Fe-binding residues (green), thiol/selenol-binding residues (red), and histidine/hercynine-binding residues (blue) are highlighted. Putative Fe-binding residues of uncharacterized subfamilies A-F have not been experimentally verified, but are highlighted green for visual clarity. See **Supplementary Fig. 2** for additional information.

| NHISS Subfamily | # of Members | Most Common Composite Motif | Example (NCBI accession, strain name) |
|---|---|---|---|
| OvoA | 4375 | RHYH--DSHIHYFYAFF | WP_092912839.1, *H. thermophila* |
| EgtB-IV | 466 | RNYH--DTHIHYQNWFF | WP_008197519.1, *M. aeruginosa* |
| EgtB-II | 16304 | ASHHYYRRHQHVYNFFA | WP_014099805.1, *C. thermophilum* |
| EgtB-I | 12211 | MSDH--RRHQHYYNWDR | EHI13262.1, *M. thermoresistibile* |
| SenA | 1514 | VNEH--HRHMHYYNFYR | WP_062361878.1, *V. paradoxus* |
| OvsA | 228 | VNEH--HRHMHYMQAYR | WP_077530365.1, *H. utahensis* |
| Unknown A | 42 | GAHHRDPTHRHDVHYLS | ALJ35470.1, *A. brasilense* |
| Unknown B | 547 | FSHH--KRHQHYYNWYV | BAC08952.1, *T. vestitus* |
| Unknown C | 73 | LNQH--HRHLCWYAFHR | MCC7716603.1, *J. lividum* |
| Unknown D | 226 | HATQ--PCHLRVLGFLR | CCF96682.1, *R. solanacearum* |
| Unknown E | 112 | ADTH--PRHAQSDWARD | AEI77914.1, *C. necator* |
| Unknown F | 75 | VVLH--PRHLRWGGFAR | WP_018903115.1, *V. paradoxus* |

**Supplementary Table 5.** NMR assignments for 5-SeHis-diselenide, collected in $D_2O$. The structure and numbering scheme are shown at the top of the table. See **Supplementary Fig. 7** for NMR spectra.

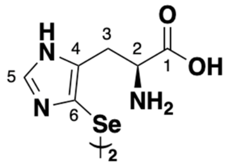

| Pos | δH (m, J (Hz)) | δC |
|---|---|---|
| 1 | - | 172.9 |
| 2 | 3.71 (dd, 7.7, 5.5) | 54.3 |
| 3 | 2.46 (dd, 15.2, 7.8); 2.56 (dd, 15.2, 5.5) | 26.2 |
| 4 | - | 138.0 |
| 5 | 7.84 (s) | 138.9 |
| 6 | - | 116.1 |

**Supplementary Table 6.** NMR assignments for ovoselenol-diselenide, collected in $D_2O$. The structure and numbering scheme are shown at the top of the table. See **Supplementary Fig. 8** for NMR spectra.

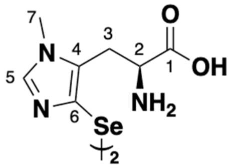

| Pos | δH (m, J (Hz)) | δC |
|---|---|---|
| 1 | - | 172.5 |
| 2 | 3.62 (t, 7.5) | 53.5 |
| 3 | 2.60 (dd, 15.3, 7.8); 2.80 (dd, 15.3, 5.0) | 24.5 |
| 4 | - | 133.0 |
| 5 | 7.69 (s) | 140.5 |
| 6 | - | 124.5 |
| 7 | 3.58 (s) | 32.0 |

**Supplementary Table 7.** Crystallographic data processing and refinement statistics for OvsA structures.

| PDB ID (Ligand) | 8U42 (Wild-type OvsA + iron) | 8U41 (Wild-type OvsA + iron + histidine) | 8UX5 (OvsA-YNF + iron) |
|---|---|---|---|
| **Data Collection**[a] | | | |
| Space group | P $2_1$ $2_1$ $2_1$ | P $6_5$ | P $6_5$ |
| Unit cell (Å) | a = 108.22, b = 115.41, c = 121.71<br>α = β = γ = 90 | a = b = 160.129, c = 122.312<br>α = β = 90, γ = 120 | a = b = 162.174, c = 124.663<br>α = β = 90, γ = 120 |
| Wavelength (Å) | 1.0332 | 0.97872 | 0.9686 |
| Resolution range (Å) | 57.71 – 2.0 (2.071 – 2.0) | 69.34 – 2.72 (2.817 – 2.72) | 29.09 – 3.06 (3.169 – 3.06) |
| Total observations | 183644 (18439) | 95745 (9480) | 70247 (6906) |
| Total unique observations | 98747 (9984) | 47873 (4740) | 35167 (3456) |
| $CC_{1/2}$ | 0.992 (0.562) | 0.999 (0.654) | 0.999 (0.501) |
| $I/\sigma_1$ | 6.57 (1.27) | 16.89 (1.50) | 14.10 (1.09) |
| Completeness (%) | 95.18 (96.39) | 99.93 (99.92) | 99.84 (99.97) |
| $R_{merge}$ | 0.07203 (0.6019) | 0.02925 (0.5307) | 0.03673 (0.6614) |
| $R_{pim}$ | 0.07203 (0.6019) | 0.02925 (0.5307) | 0.03673 (0.6614) |
| Redundancy | 1.9 (1.8) | 2.0 (2.0) | 2.0 (2.0) |
| **Refinement Statistics** | | | |
| Resolution range (Å) | 57.71 – 2.0 | 69.34 – 2.72 | 29.09 – 3.06 |
| Reflections (total) | 98353 (9836) | 47843 (4736) | 35154 (3455) |
| Reflections (test) | 1997 (195) | 1967 (199) | 1752 (173) |
| Total atoms refined | 7963 | 7296 | 7055 |
| Solvent | 712 | 85 | 30 |
| $R_{work}$ ($R_{free}$) | 0.1832 (0.2087) | 0.2247 (0.2462) | 0.2029 (0.2530) |
| RMSDs | | | |
| Bond lengths (Å)/angles (°) | 0.014/1.22 | 0.005/0.88 | 0.008/1.10 |
| Ramachandran plot | | | |
| Favored/allowed (%) | 96.84/3.16 | 92.30/7.03 | 93.17/6.38 |
| Mean B values (Å$^2$) | | | |
| Protein | 35.23 | 78.76 | 94.54 |
| Ligands | 42.42 | 69.57 | 75.88 |
| Solvent | 41.13 | 65.75 | 77.67 |

[a] Values in parentheses refer to the high-resolution shell.

**Supplementary Table 8.** Top ranking structures (ranked by the RMSD) homologous to OvsA, as identified by the Dali protein structure comparison server.

| PDB accession code | Chain | RMSD Cα [Å] | Dali Z-score | % Identity | Description (donor organism) | Ref |
|---|---|---|---|---|---|---|
| 8K5J | A | 1.8 | 41.4 | 35 | Selenoxide synthase SenA (*Variovorax paradoxus*) | (15) |
| 4X8B | B | 2.2 | 47.3 | 29 | Sulfoxide synthase EgtB-I (*Mycolicibacterium thermoresistibile*) | (13) |
| 2Q17 | B | 2.2 | 21.2 | 30 | Formylglycine-generating enzyme (*Streptomyces coelicolor*) | (50) |
| 6O6L | D | 2.3 | 41.7 | 29 | Sulfoxide synthase EgtB-II (*Chloracidobacterium thermophilum*) | (51) |
| 5HHA | A | 2.4 | 19.8 | 25 | Periplasmic oxidoreductase PvdO (*Pseudomonas aeruginosa*) | N/A |
| 5NYY | A | 2.5 | 21.3 | 33 | Formylglycine-generating enzyme (*Thermomonospora curvata*) | (52) |
| 2Y3C | A | 2.5 | 19.4 | 21 | Diversity-generating retroelement variable protein A (*Treponema denticola*) | (53) |
| 2IOU | D | 2.5 | 12.6 | 15 | Major tropism determinant P1 (*Bordetella brochiseptica*) | (54) |
| 2HIB | A | 2.6 | 21.4 | 31 | Formylglycine-generating enzyme (*Homo sapiens*) | (55) |
| 8KHQ | A | 2.7 | 36.1 | 25 | Sulfoxide synthase OvoA (*Hydrogenimonas thermophila*) | (14) |
| 5AOH | B | 3.1 | 18.3 | 22 | CarF (*Serratia sp. ATCC 39006*) | (56) |

**Supplementary Figure 1.** Detailed biosynthetic pathways of TSIs[4,8-11,27]. (a,b) Biosynthesis of thiol/selenol substrates of NHISS enzymes. (c) Biosynthesis of ovothiol and ergothioneine, notably requiring a enzyme-catalyzed C–S bond cleavage step by OvoB or EgtE, respectively. (d) Biosynthesis of selenoneine and ovoselenol, featuring a spontaneous selenoxide elimination resulting in C–Se bond cleavage. (e) Qualitative cladogram of characterized NHISS enzyme sequences. (f) The standalone methyltransferase OvsM shares strong sequence similarity with the *C*-terminal methyltransferase domain of OvoA[8,14,27], despite low similarity between OvsA and the *N*-terminal NHISS domain of OvoA. (g) Average pairwise sequence similarity between members of characterized NHISS subfamilies, respresented as -log(E-value).

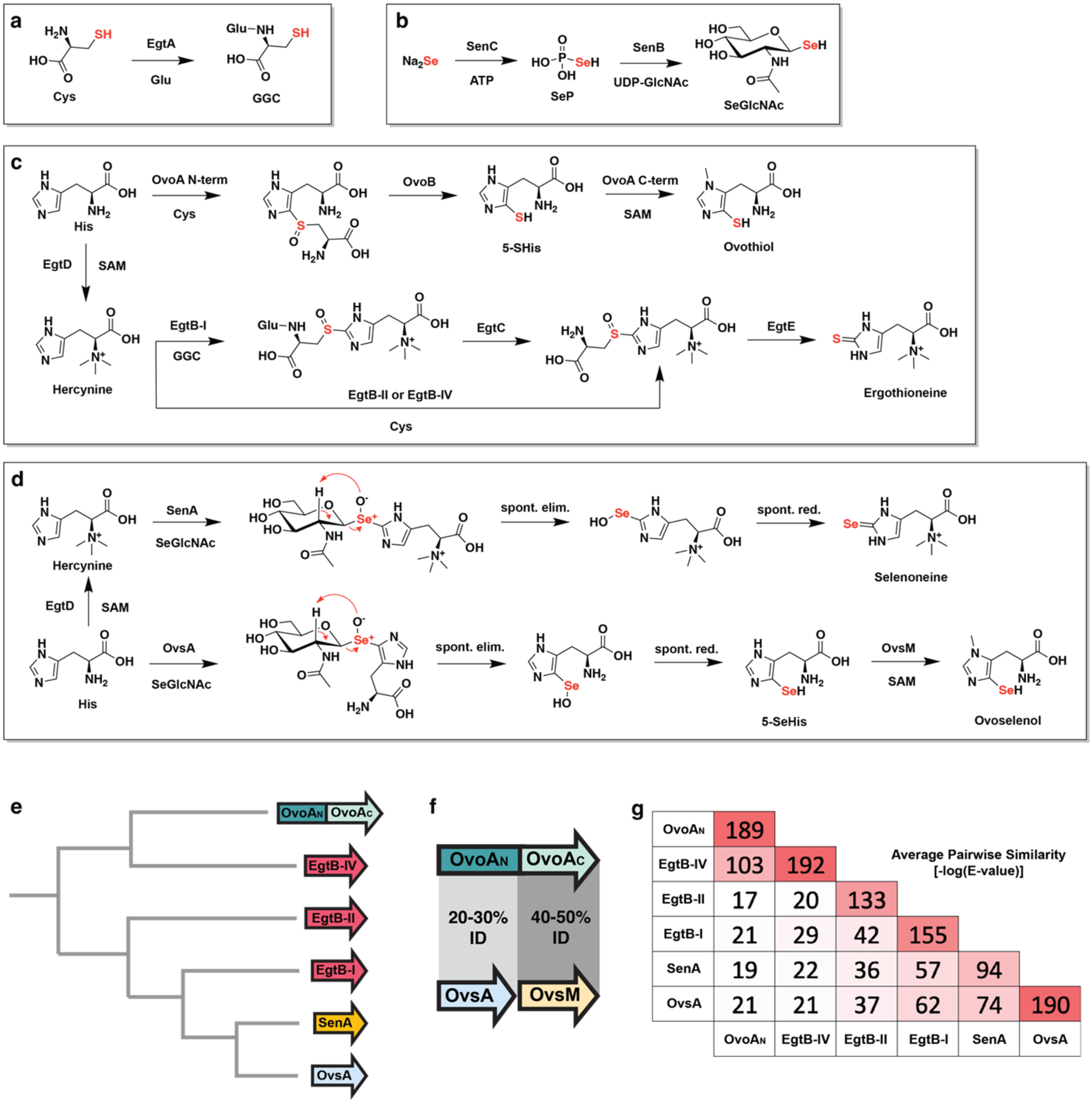


| | OvoAN | EgtB-IV | EgtB-II | EgtB-I | SenA | OvsA |
|---|---|---|---|---|---|---|
| OvoAN | 189 | | | | | |
| EgtB-IV | 103 | 192 | | | | |
| EgtB-II | 17 | 20 | 133 | | | |
| EgtB-I | 21 | 29 | 42 | 155 | | |
| SenA | 19 | 22 | 36 | 57 | 94 | |
| OvsA | 21 | 21 | 37 | 62 | 74 | 190 |

**Supplementary Figure 2.** Phylogenetic tree of NHISS enzymes (**a**), expanded to include representatives from seven additional uncharacterized subfamilies that feature unique composite motifs (**b**). Putative Fe-binding residues of uncharacterized subfamilies have not been experimentally verified, but are highlighted green for visual clarity. See **Supplementary Table 4** for additional information.

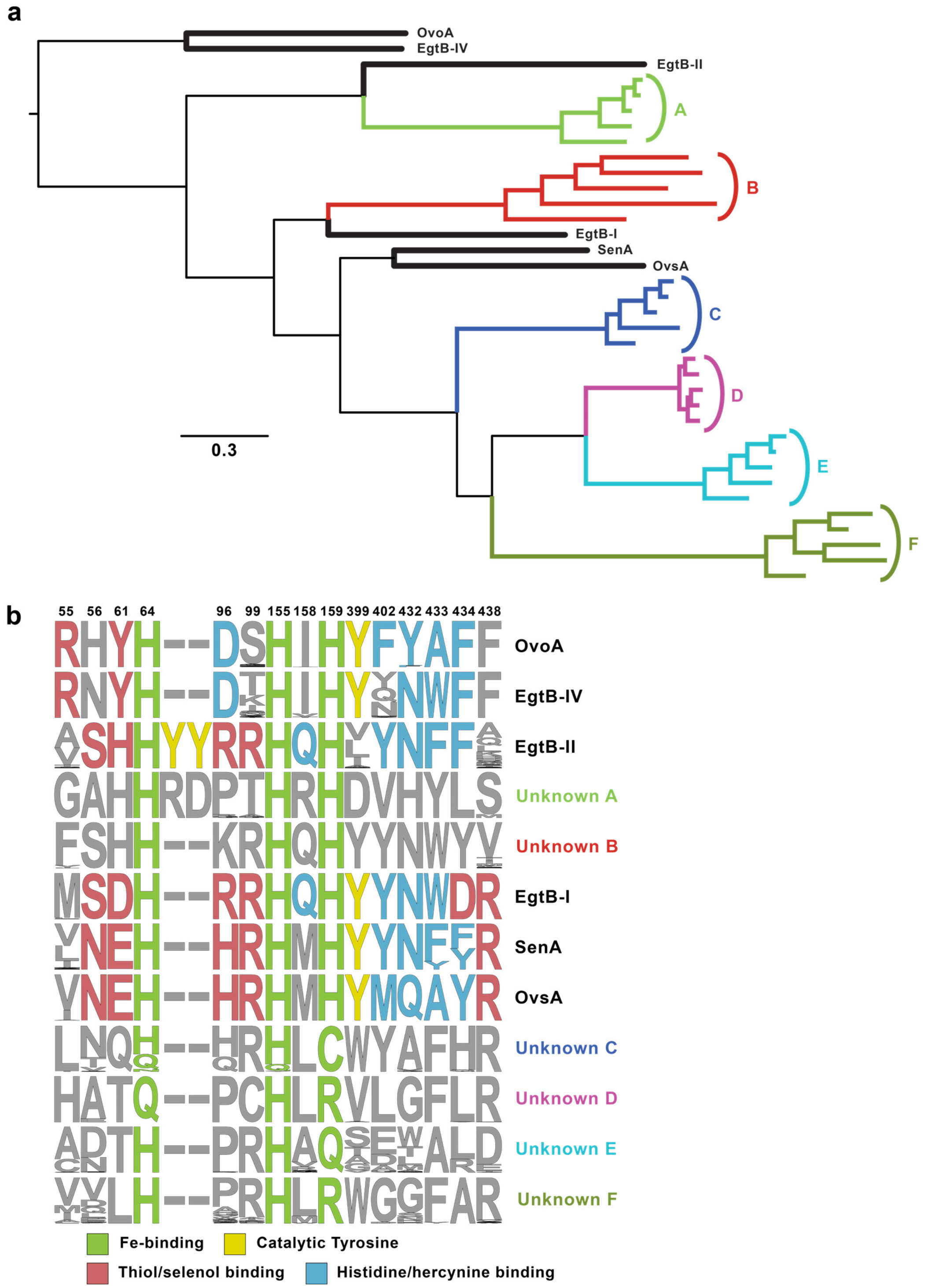

**Supplementary Figure 3.** *Ovs* cluster encoded in microbial genomes. (**a**) Representative *ovs* clusters. Red hash marks indicate chromosomal separation of greater than 10 kb. (**b**) *Limnobacter thiooxidans* represents a rare example of an organism that encodes the biosynthesis of all four TSIs.

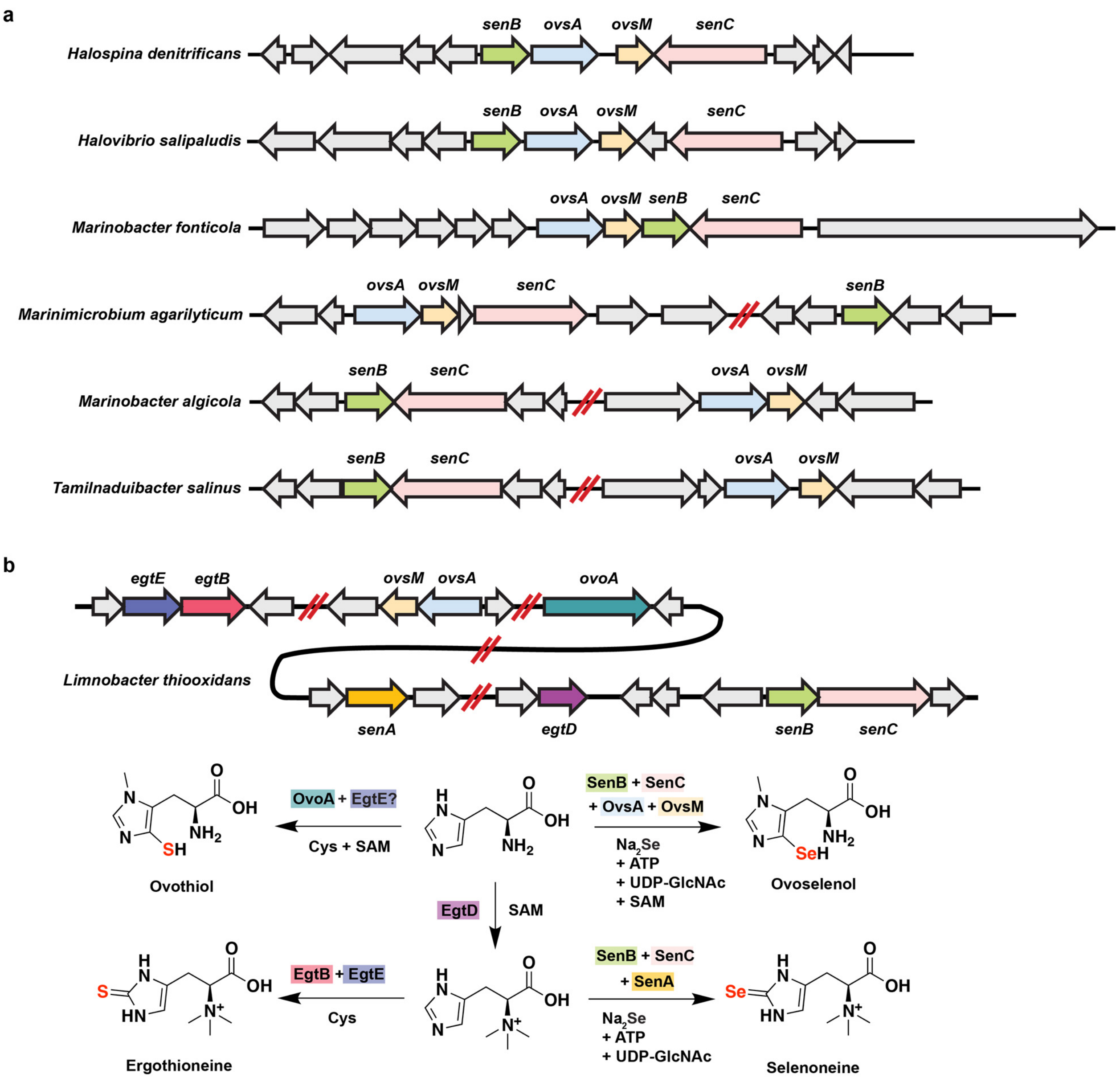

**Supplementary Figure 4.** Order of reactions in the *ovs* pathway. (**a**) OvsA fails to selenylate Nπ-methylhistidine (Nπ-MeHis) and OvsM fails to methylate histidine, suggesting a clear order of biosynthetic events. (**b**) Extracted ion chromatograms of His, Nπ-MeHis, 5-SeHis-diselenide, and OVS-diselenide (all underivatized) from OvsA/M reactions with various substrates.

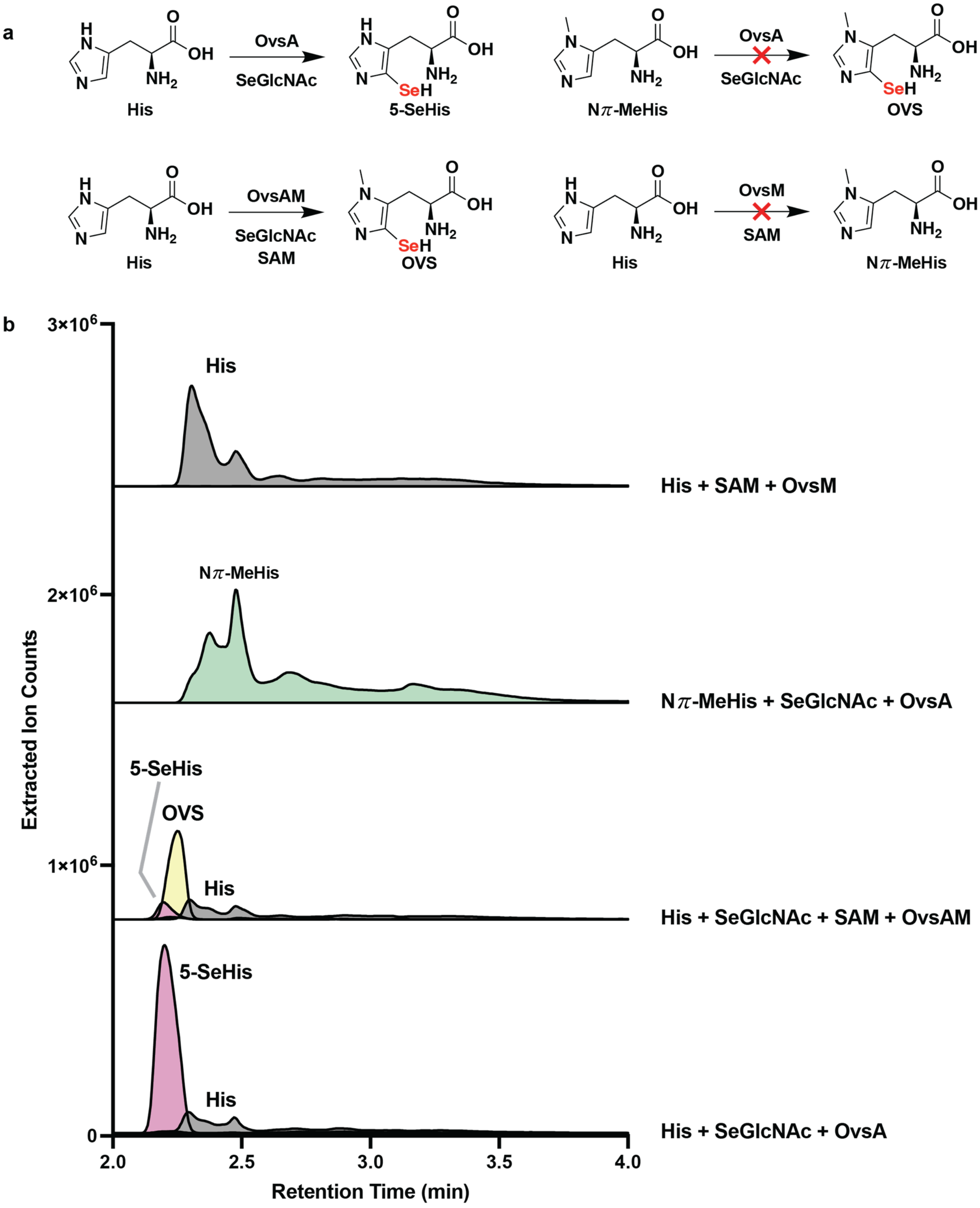

**Supplementary Figure 5.** Detection of 2-acetamidoglucal (2-AG) as a product of the OvsA reaction. (**a**) 2-AG is generated by spontaneous selenoxide elimination, following oxidative C–Se bond formation by OvsA. (**b**) 2-AG is detected only in the presence of histidine, SeGlcNAc, and OvsA. Omission of OvsA, or replacement of histidine with a non-substrate histidine derivative, yields trace or no 2-AG. Extracted ion chromatograms of 2-AG ($[M+Na]^+$ = 226.0686) are shown.

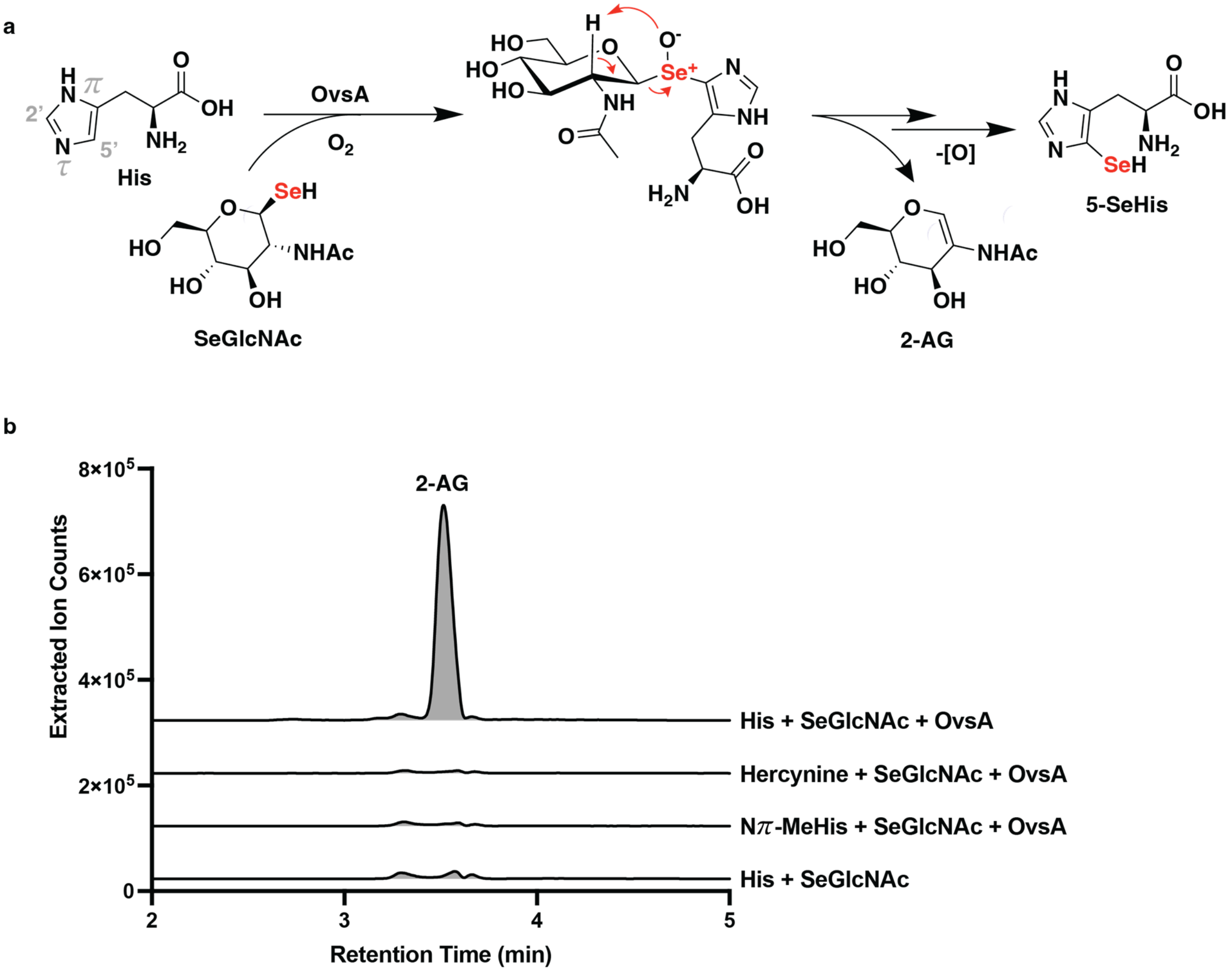

**Supplementary Figure 6.** PCR verification of *M. vinifirmus* Δ*ovsA* genotype. Red arrows indicate PCR primers used to amplify the chromosomal region of the Δ*ovsA* strain that contains the $Cm^R$ insertion.

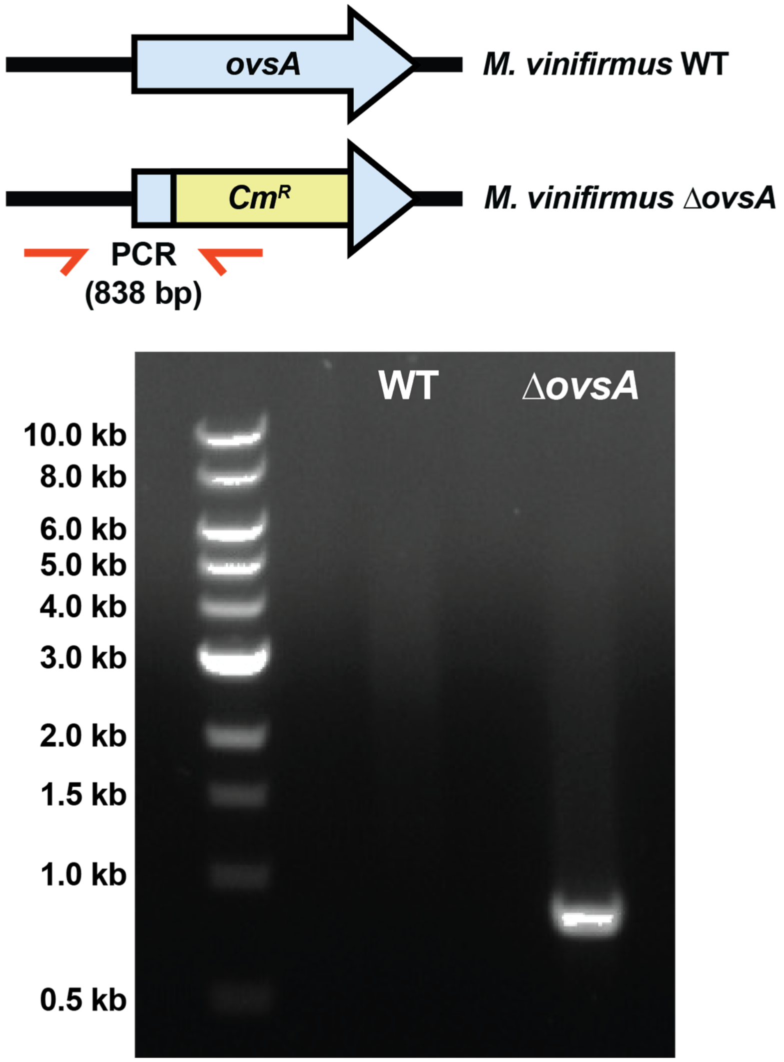

**Supplementary Figure 7.** NMR spectra of 5-SeHis-diselenide in $D_2O$ at 500 MHz on pages **S17-S20**. Shown are from top to bottom $^1H$, $^1H$-$^1H$ COSY, $^1H$-$^{13}C$ HSQC, and $^1H$-$^{13}C$ HMBC spectra.

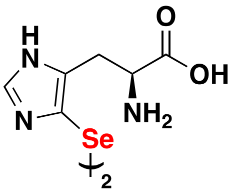


**5-SeHis-diselenide**

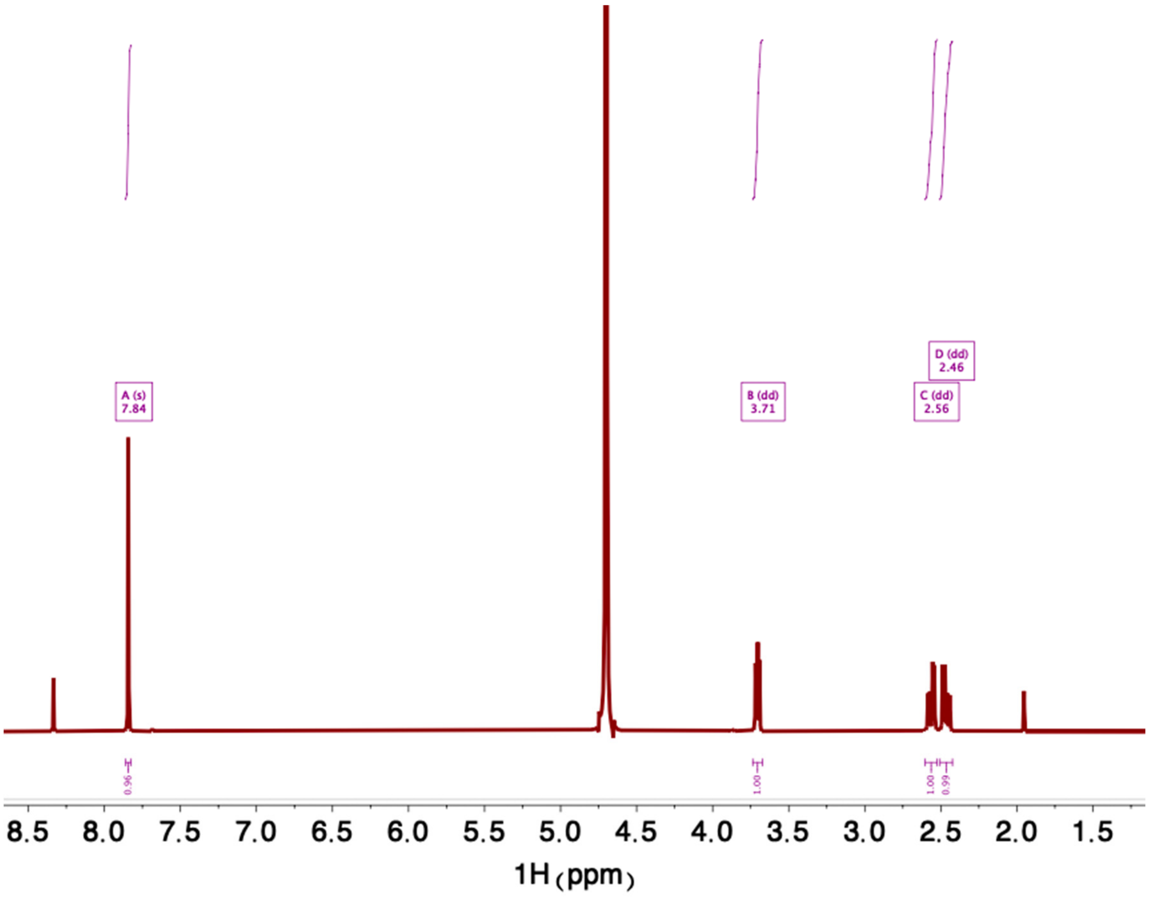

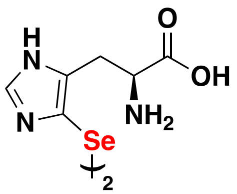

**5-SeHis-diselenide**

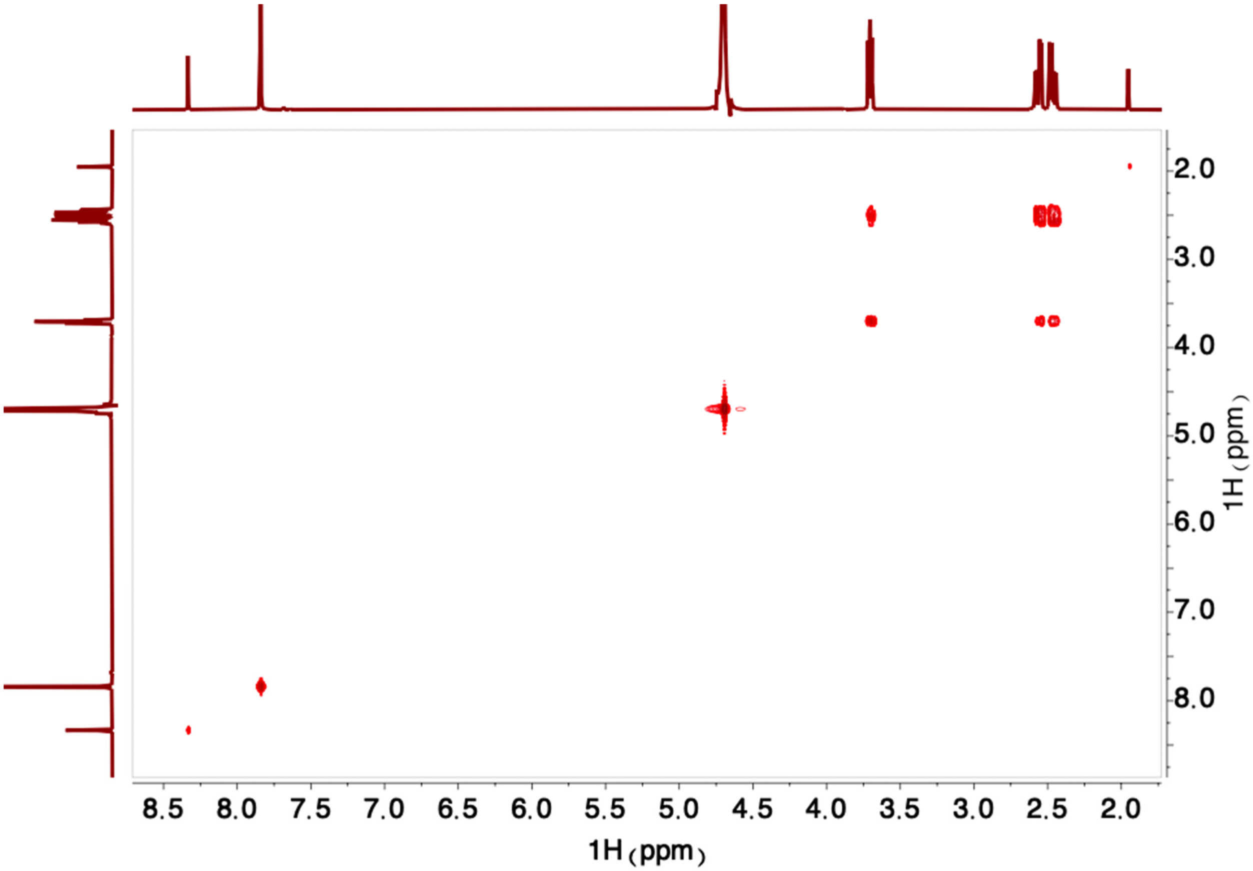

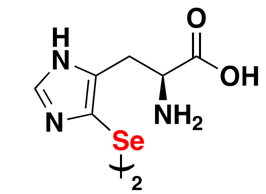


**5-SeHis-diselenide**

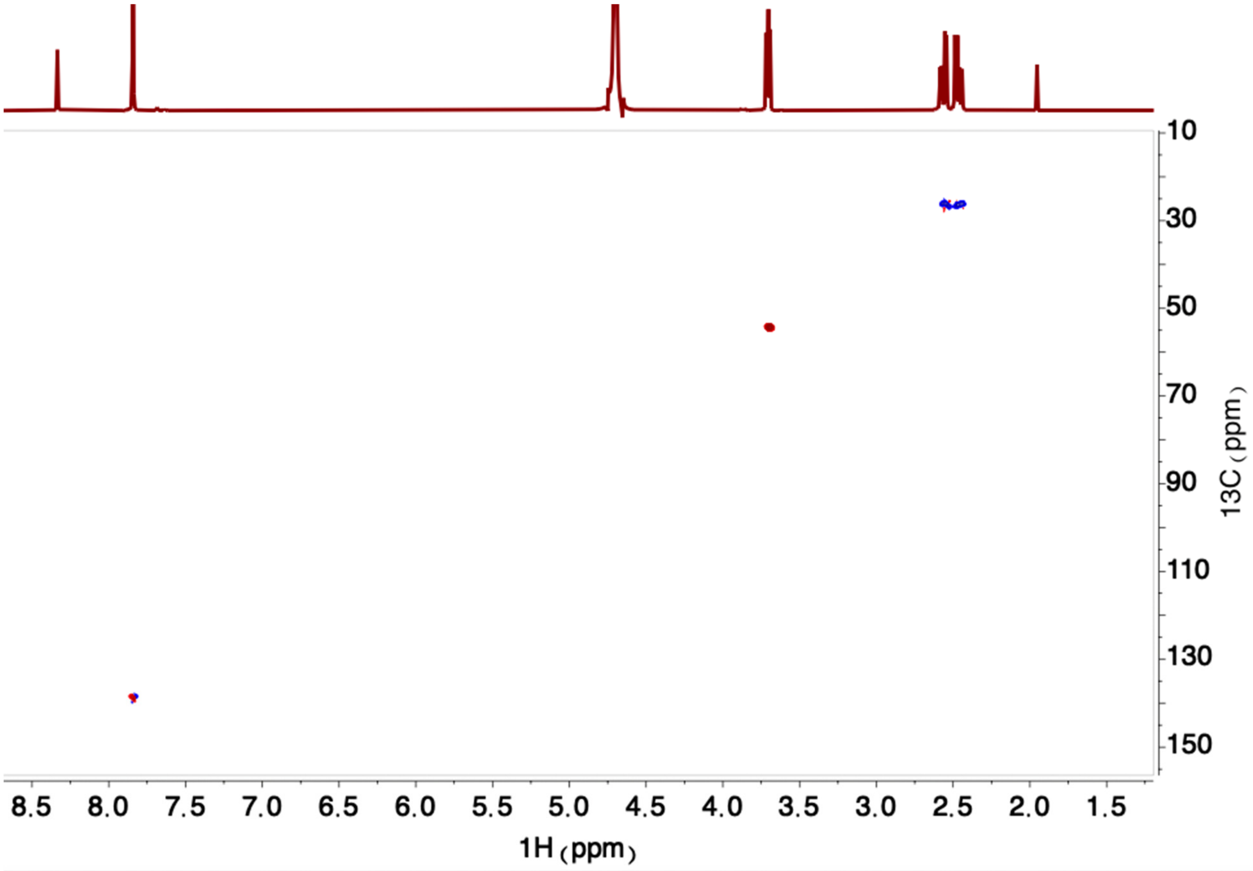

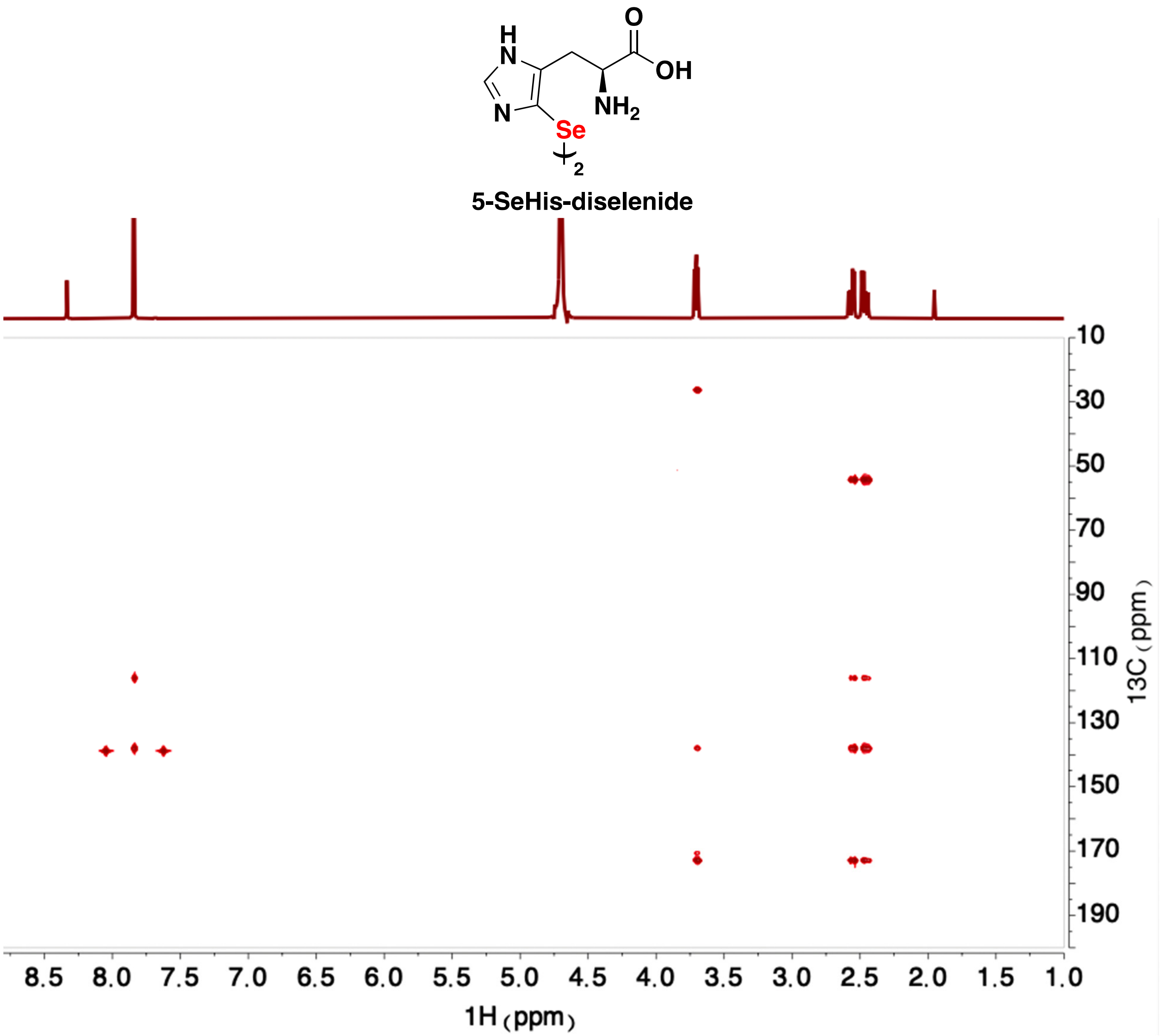

H
N
N
Se
2
O
OH
NH2
5-SeHis-diselenide
8.5
8.0
7.5
7.0
6.5
6.0
5.5
5.0
4.5
4.0
3.5
3.0
2.5
2.0
1.5
1.0
1H (ppm)
10
30
50
70
90
110
130
150
170
190
13C (ppm)

**Supplementary Figure 8.** NMR spectra of ovoselenol-diselenide in $D_2O$ at 500 MHz on pages **S21-S23**. Shown are from top to bottom $^{1}H$, $^{1}H$-$^{13}C$ HSQC, and $^{1}H$-$^{13}C$ HMBC spectra.

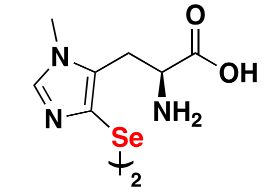


**Ovoselenol-diselenide**

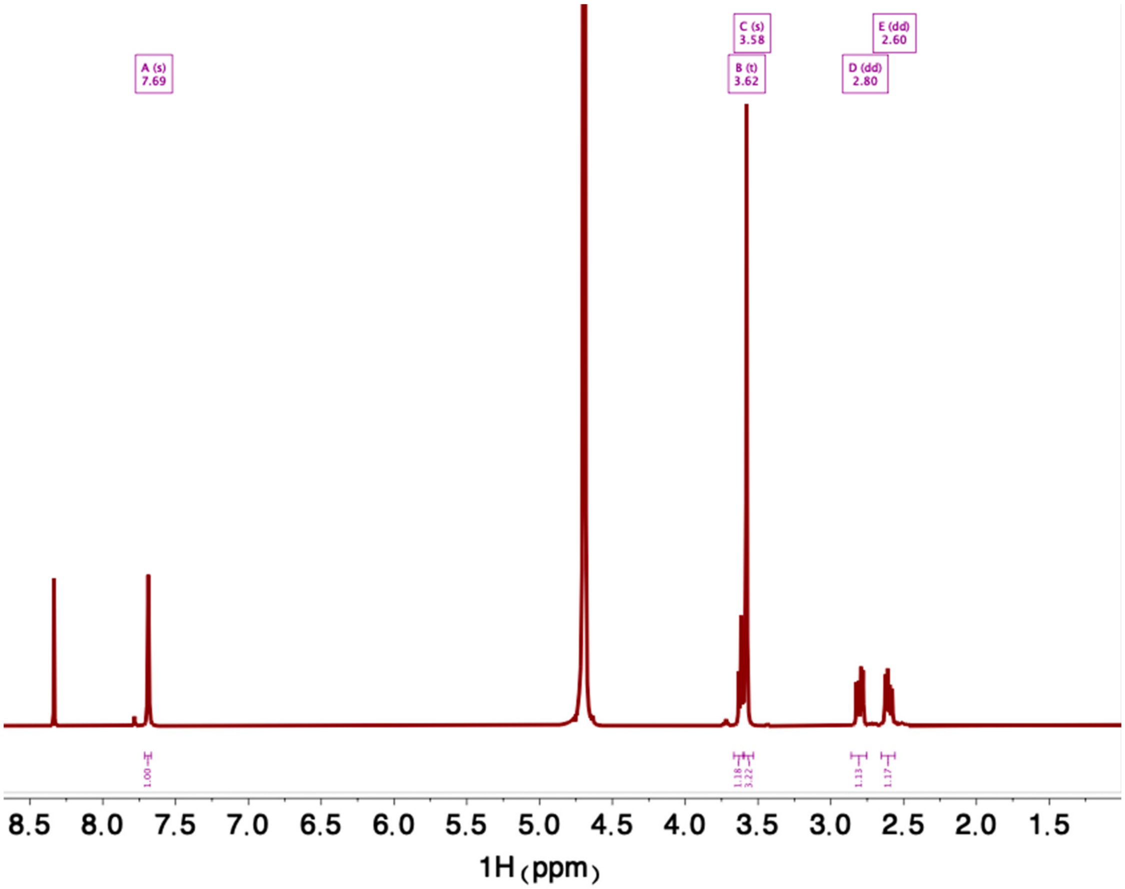

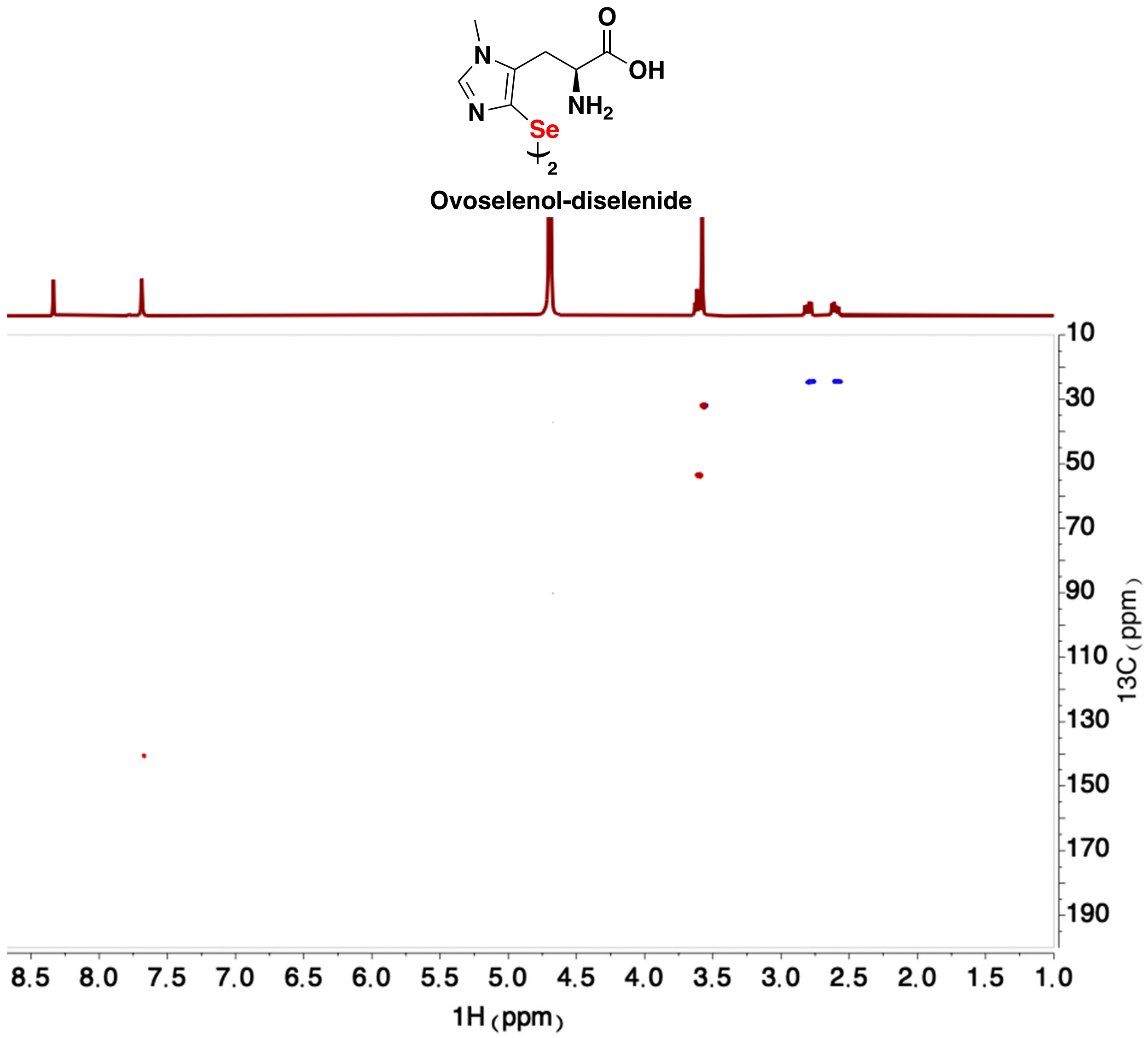
O
OH
N
N
NH2
Se
2
Ovoselenol-diselenide
10
30
50
70
90
110
130
150
170
190
13C (ppm)
8.5
8.0
7.5
7.0
6.5
6.0
5.5
5.0
4.5
4.0
3.5
3.0
2.5
2.0
1.5
1.0
1H (ppm)

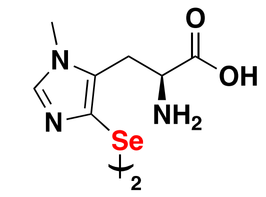


**Ovoselenol-diselenide**

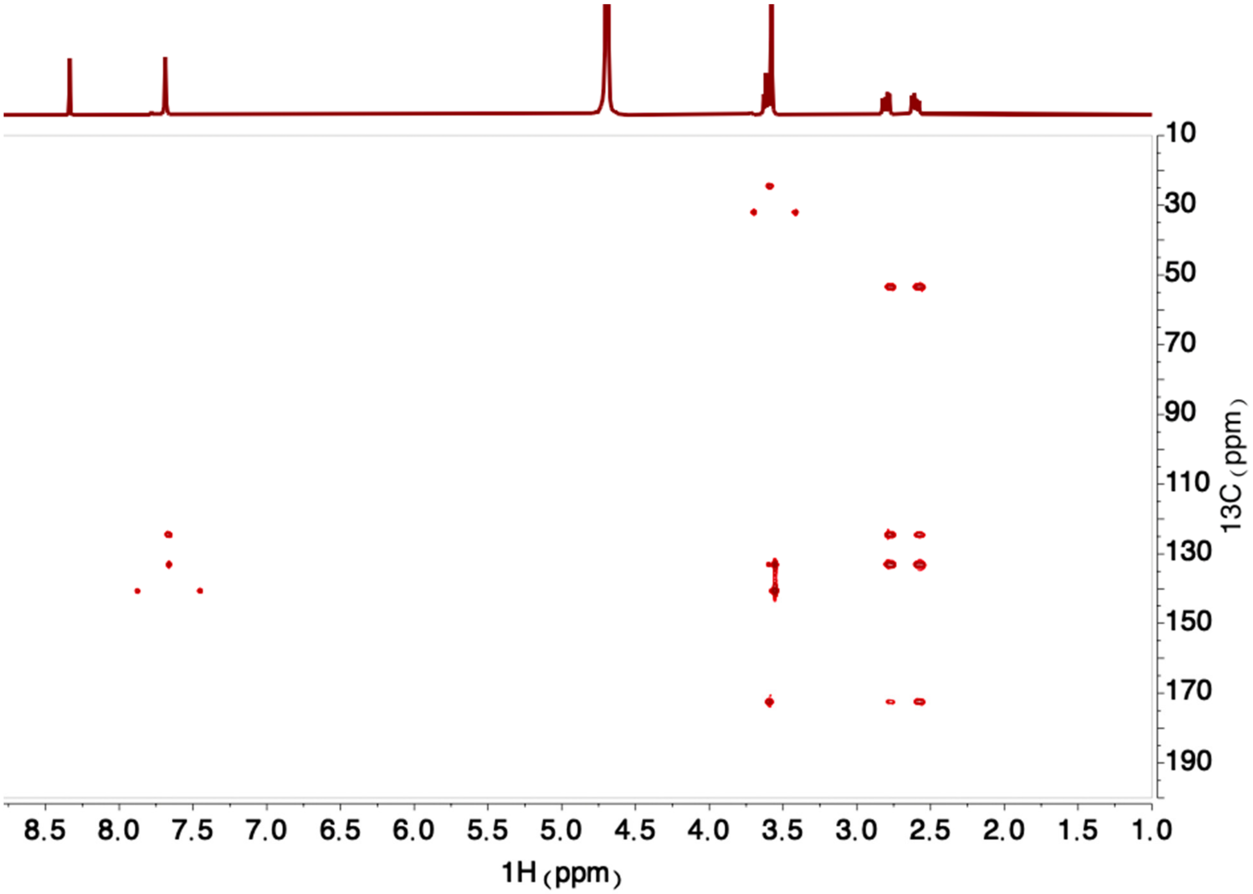

### Additional Discussion of OvsA Structure

#### OvsA's *N*-terminal domain adopts a DinB-like fold

OvsA's *N*-terminal DinB domain (residues 1-146) is characterized by a four-helix bundle with an atypical down-up-up-down topology[20], due to the long linker between α-helices 2 and 3, which forms two additional minor α-helices (α2′ and α2′′). This unusual fold has been previously reported for other proteins in the DinB superfamily, namely YfiT from *Bacillus subtilis*[57], the mycothiol-dependent maleylpyruvate isomerase from *Corynebacterium glutamicum*[58], and TTHA0303 from *Thermus thermophilus*[59]. Helices 1 and 2 are also connected by a long loop which forms a third minor α-helix (α1′). The four-helix bundle has a hydrophobic core composed primarily of Leu, Ile, Val, and Met residues. Helices 2 and 4 contain the His residues (47, 130, and 134) responsible for coordinating the mononuclear iron cofactor with octahedral geometry. This three-His facial triad is conserved for several members of the DinB superfamily, most of which coordinate a nickel cofactor rather than iron[20].

#### OvsA's *C*-terminal domain adopts a C-type lectin fold

The *C*-terminal CLec domain (residues 147-448) hosts many flexible loop regions and few secondary structure elements, containing seven short α-helices and four antiparallel β-sheets (nine total β-strands). In addition to the other structurally characterized NHISS enzymes, structural homology searches using the Dali server[60] identified several other structural homologs of OvsA containing the loop-rich CLec fold, including the FGEs (RMSD of 2.6 Å for the human enzyme, PDB accession code 2HIB)[50,52,55], the periplasmic oxidoreductase PvdO from *Pseudomonas aeruginosa* (RMSD of 2.4 Å, PDB accession code 5HHA), and the diversity-generating retroelement variable protein A (TvpA) from *Treponema denticola* (RMSD of 2.5 Å, PDB accession code 2Y3C)[53] (**Supplementary Table 8**). In OvsA, this loop-rich domain is stabilized by an extensive network of hydrogen bonds and ionic interactions between Arg and Glu/Asp residues. A sodium ion is coordinated at the interior of the CLec domain, by the backbone carbonyls of residues Gln374, Met375, Gly377, Val379, and the side chain of Glu381. The bound cation is present in the same location for all OvsA structures described herein as well as EgtB-I, the FGEs, PvdO, and other CLec proteins[53,56], in the form of a sodium, potassium, or calcium ion. We suspect that the bound ion provides stabilization for this loop-rich region rather than participating in catalysis, as evidenced by its distant location from the active site.

**Supplementary Figure 9.** Analysis of TSIs by cyclic voltammetry (CV) and linear sweep voltammetry (LSV). (**a**) CV analysis of ovoselenol (black trace) shows that the reaction is non-reversible and complex. Because ovoselenol is isolated in the diselenide form, it can be first electrochemically reduced by holding the potential at -0.55 V (vs. SCE), followed by a linear oxidative sweep, as shown by the LSV trace (red). The peak oxidation potential observed by CV and LSV is similar and is consistent with an analogous oxidation reaction in both cases. The small shift in peak potential is likely due to minor concentration differences in the two samples. (**b**) Scheme for the CV/LSV experiments. The reduction reaction likely involves an electron transfer–diselenide/sulfide bond cleavage–electron transfer process, as determined for other disulfides[19]. In the case of the diselenide form of ovoselenol, once reduction occurs, the anionic monomeric form accumulates, due to the highly acidic nature of the selenol group, and one-electron oxidation may be observed, as shown, in the oxidative sweep. Note that the peak potentials here and in **Fig. 3i** do not represent the standard thermodynamic reduction/oxidation potential, but rather provide a qualitative and relative measure of the antioxidant capacity of the four molecules under examination.

**a**

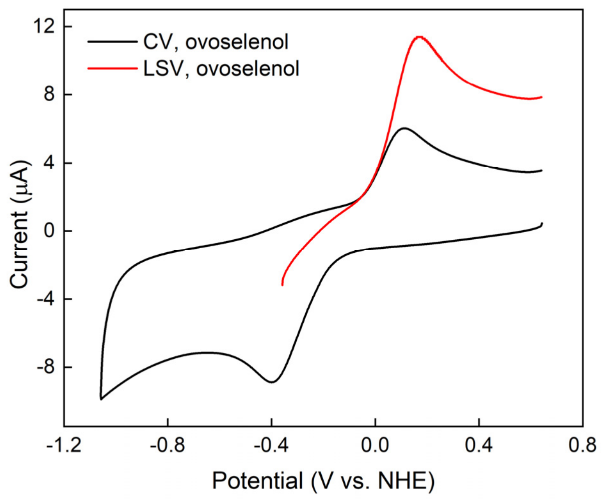


**b**

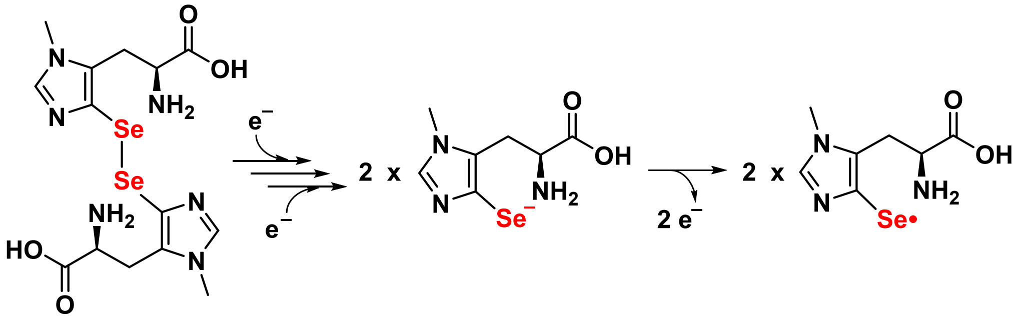

**Supplementary Figure 10.** Structural alignment of OvsA (light blue, PDB accession code 8U42) and its closest structural homolog, SenA (light gray, PDB accession code 8K5K). The tertiary structures align very closely, with small differences in loops vs. secondary structure elements. Secondary structure elements present only in OvsA are shown in dark blue while secondary structure elements present only in SenA are shown in dark gray. The primary differences are the presence of an additional loop in OvsA (residues 250-265, black arrow) and a longer loop region connecting α-helices 2 and 3 in SenA (residues 81-97, dark pink arrow).

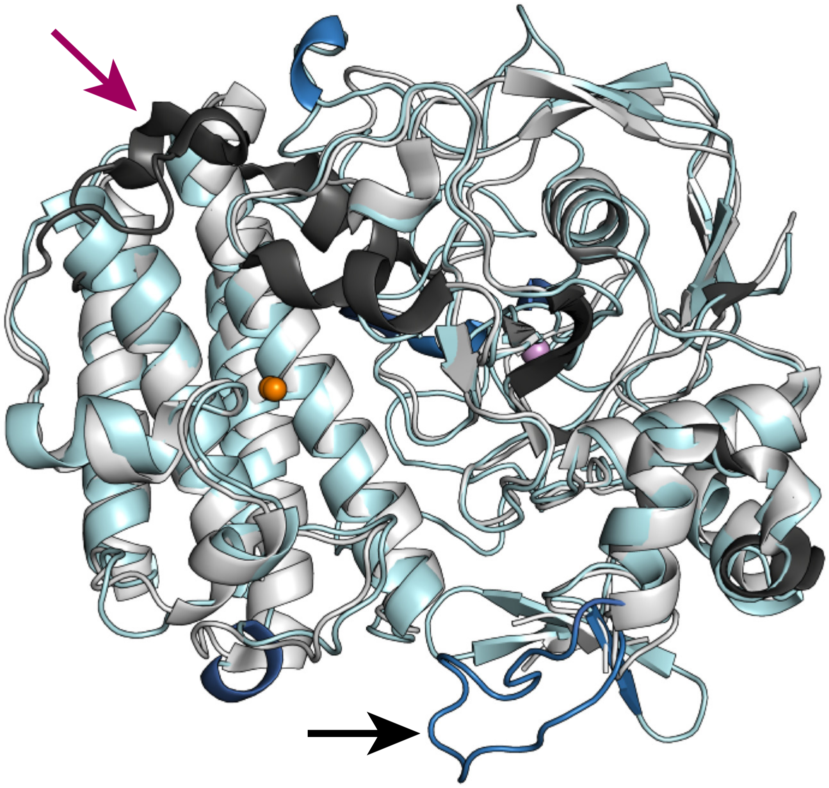

**Supplementary Figure 11.** Polder map ($F_o$-$F_c$) contoured at 7.0 sigma for iron-bound histidine in OvsA, viewed from the front (a) and from the side (b).

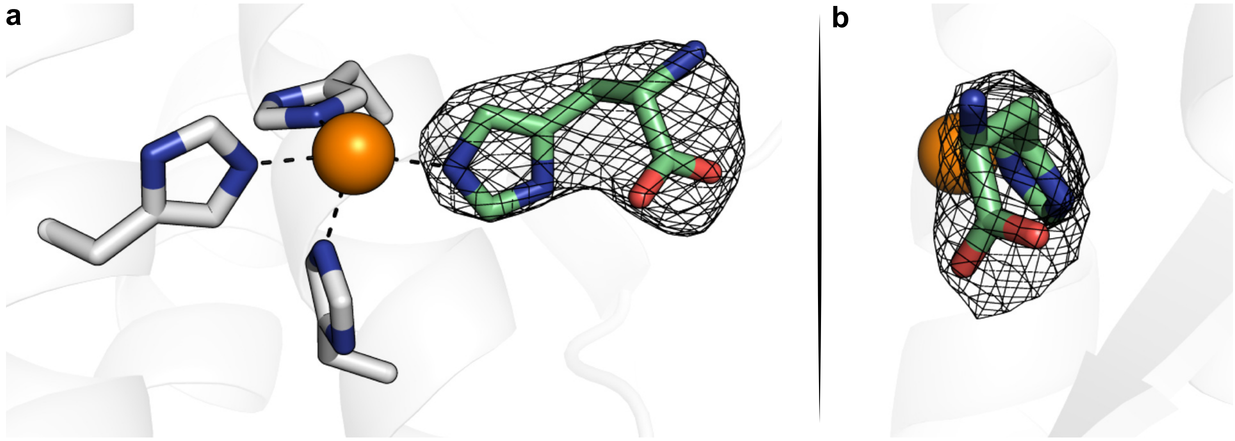

**Supplementary Figure 12.** Overlaid active sites of OvsA (green), OvsA-YNF (dark blue), and SenA (pale blue). In OvsA-YNF, Gln405 rotates to form two additional water-mediated hydrogen bonds to the imidazole π-nitrogen of hercynine through the same water molecule involved in the hydrogen bond with the Y401 backbone carbonyl. A Trp residue is present at the same location in SenA (W370), which forms a slightly longer hydrogen bond to this same water molecule.

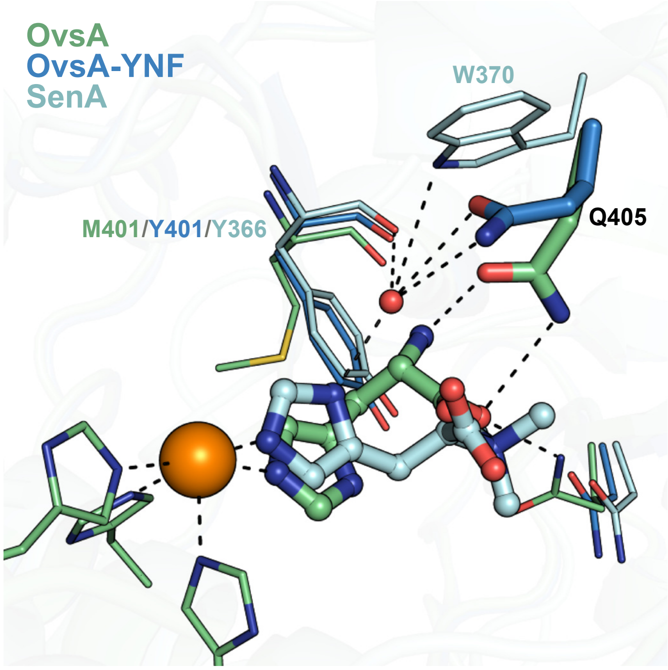

### Supplementary References


48. Sorokin, D. Yu., & Tindall, B. J. The status of the genus name Halovibrio Fendrich 1989 and the identity of the strains *Pseudomonas halophila* DSM 3050 and *Halomonas variabilis* DSM 3051. Request for an opinion. *Int. J. Syst. Evol. Microbiol.* **56**, 487–489 (2006).
49. Tindall, B. J. The designated type strain of *Pseudomonas halophila* Fendrich 1989 is DSM 3051, the designated type strain of *Halovibrio variabilis* Fendrich 1989 is DSM 3050, the new name *Halomonas utahensis* (Fendrich 1989) Sorokin and Tindall 2006 is created for the species represented by DSM 3051 when treated as a member of the genus *Halomonas*... *Int. J. Syst. Evol. Microbiol.* **64**, 3588–3589 (2014).
50. Carlson, B. L. et al. Function and structure of a prokaryotic formylglycine-generating enzyme. *J. Biol. Chem.* **283**, 20117–20125 (2008).
51. Naowarojna, N. et al. Crystal structure of the ergothioneine sulfoxide synthase from candidatus *Chloracidobacterium thermophilum* and structure-guided engineering to modulate its substrate selectivity. *ACS Catal.* **9**, 6955–6961 (2019).
52. Meury, M., Knop, M. & Seebeck, F. P. Structural basis for copper-oxygen mediated C-H bond activation by the formylglycine-generating enzyme. *Angew. Chem. Int. Ed. Engl.* **56**, 8115–8119 (2017).
53. Le Coq, J. & Ghosh, P. Conservation of the C-type lectin fold for massive sequence variation in a Treponema diversity-generating retroelement. *Proc. Natl. Acad. Sci. U.S.A.* **108**, 14649–14653 (2011).
54. Miller, J. L. et al. Selective ligand recognition by a diversity-generating retroelement variable protein. *PLoS Biol.* **6**, 1195–1207 (2008).
55. Roeser, D., Schmidt, B., Preusser-Kunze, A. & Rudolph, M. G. Probing the oxygen-binding site of the human formylglycine-generating enzyme using halide ions. *Acta Crystallogr. D Biol. Crystallogr.* **63**, 621–627 (2007).
56. Tichy, E. M., Hardwick, S. W., Luisi, B. F. & Salmond, G. P. C. 1.8 A resolution crystal structure of the carbapenem intrinsic resistance protein CarF. *Acta. Crystallogr. D Struct. Biol.* **73**, 549–556 (2017).
57. Rajan, S. S., Yang, X., Shuvalova, L., Collart, F. & Anderson, W. F. YfiT from Bacillus subtilis is a probable metal-dependent hydrolase with an unusual four-helix bundle topology. *Biochem.* **43**, 15472–15479 (2004).
58. Wang, R. et al. Crystal structures and site-directed mutagenesis of a mycothiol-dependent enzyme reveal a novel folding and molecular basis for mycothiol-mediated maleylpyruvate isomerization. *J. Biol. Chem.* **282**, 16288–16294 (2007).
59. Nagata, K. et al. Crystal structure of TTHA0303 (TT2238), a four-helix bundle protein with an exposed histidine triad from *Thermus thermophilus* HB8 at 2.0 A. *Proteins* **70**, 1103–1107 (2008).
60. Holm, L. Dali server: structural unification of protein families. *Nucleic Acids Res.* **50**, 210–215 (2022).
